# A comparison of eDNA to camera trapping for assessment of terrestrial mammal diversity

**DOI:** 10.1101/634022

**Authors:** Kevin Leempoel, Trevor Hebert, Elizabeth A. Hadly

## Abstract

Environmental DNA (eDNA) is one of the most promising approaches to meet the demand for the fast and frequent monitoring of ecosystems needed to tackle the current decline in biodiversity. However, before eDNA can establish itself as a robust alternative for mammal monitoring, comparison with existing approaches is necessary, yet has not been done. Moreover, much is unknown regarding the nature, spread and persistence of DNA shed by animals into terrestrial environments, or the optimal experimental design for understanding these potential biases.

To address some of these challenges, we compared the detection of terrestrial mammals using eDNA analysis of soil samples against confirmed species observations from a long-term (∼9-yr) camera trapping study. At the same time, we considered multiple experimental parameters, including two sampling designs, two DNA extraction kits and two metabarcodes of different sizes.

All mammals consistently recorded with cameras were detected in eDNA. In addition, eDNA reported many small mammals not recorded by camera traps, but whose presence in the study area is otherwise documented. A long metabarcode (≈220bp) offering a high taxonomic resolution, achieved a similar efficiency as a shorter one (≈70bp) and a phosphate buffer-based extraction gave similar results as a total DNA extraction method for a fraction of the price. Our results support that eDNA-based monitoring should become a valuable part of terrestrial mammal surveys. Yet, the lack of coverage of mammal mitochondrial genomes in public databases must be addressed before eDNA can be used to its full potential.

## Introduction

Biodiversity loss due to human activities has been documented by numerous studies and calls for improved evaluations of species diversity, distribution and abundance (Barnosky et al., 2012; Mendenhall, Karp, Meyer, Hadly, & Daily, 2014). Ideally, frequent surveys would be used to obtain an unbiased evaluation of faunal diversity or to document changes over time. But faunal diversity surveys remain time-consuming, expensive, invasive and usually limited in terms of taxa covered (Deiner, Bik, et al., 2017; Stem, Margoluis, Salafsky, & Brown, 2005). Moreover, distinguishing cryptic species, i.e. species that look identical but that are genetically distinct, remains an arduous task as surveys are based either on photographs or measurements of phenotypic traits, requiring substantial expertise and comprehensive sampling (Bickford et al., 2007). For these reasons, biodiversity surveys remain difficult to perform despite being crucial for the successful implementation of large-scale conservation efforts (Burton et al., 2015; Frishkoff et al., 2014; Mendenhall, Shields-Estrada, Krishnaswami, & Daily, 2016).

For terrestrial mammals in particular, camera trapping is an increasingly utilized approach but needs long temporal coverage, remains expensive, demands substantial maintenance and generally does not detect small animals as large animals are usually targeted (Burton et al., 2015; O’Connell, Nichols, & Karanth, 2010). In addition, post-processing remains laborious as manual tagging of images is required, although promising developments with machine learning have been made recently (Tabak et al., 2018). For small mammals, live-trapping is more common but the type of trap, bait and sampling design can strongly affect the probability of detection of particular species (Bovendorp, McCleery, & Galetti, 2017; Harkins, Keinath, & Ben-David, 2019). Moreover, trapping is highly invasive, labor-intensive and generally requires onerous permitting.

Terrestrial environmental DNA (eDNA) is poised to become an effective alternative to existing monitoring approaches (Deiner, Bik, et al., 2017). For animals, the premise of eDNA is that pieces of skin, fur, feces or saliva are shed in the environment and that, by collecting environmental samples such as water or soil, we should be able to identify to which species the extracted DNA belongs. As such, the presence of a large panel of taxa could be evaluated non-invasively by a single approach within a given environmental sample (Drummond et al., 2015). Yet a comprehensive comparison of species diversity identified through eDNA alongside traditional surveys is required prior to using eDNA for biodiversity monitoring across ecosystems (Deiner, Bik, et al., 2017). A small number of such studies have been conducted in aquatic ecosystems and have reported a higher sensitivity of eDNA combined with a lower sampling effort when compared to visual surveys (Port et al., 2016; Thomsen et al., 2012; Valentini et al., 2016). Only a few studies have aimed to evaluate eDNA’s potential for the detection of terrestrial mammals (Andersen et al., 2012; Ishige et al., 2017; Ushio et al., 2017) and these were primarily proof-of-concept studies, mostly done in enclosed environments (e.g. fenced reserve, zoo), which may not be directly transferrable to interpreting whether eDNA from environmental samples in natural environments can reflect the local present-day mammal communities.

Many questions remain unanswered about the potential of eDNA in natural environments. For example, we do not know how frequently an animal must pass by a given area to be detectable in an eDNA sample, or how recent that passage must be. The size and behavior of an animal likely affects the amount of DNA it leaves in the environment (Adams, Hoekstra, Muell, & Janzen, 2019), meaning that some animals may only rarely be sampled while others may be over-represented. On the methodological side, unanswered questions include the volume and number of environmental samples that should be collected, which environmental source is the most versatile, and most importantly, whether all target species are detectable and if eDNA reflects their abundance. Previous studies have provided partial answers to these questions. For example, Andersen *et al.* (2012) found that the detection rate was higher when combining subsamples in a large grid than from one unique point, but the amount of soil they collected per site was low (6.5g). Their study showed that top soil is a relevant source of mammal DNA, with the advantage that it is unlikely to move over long distances, in contrast to aquatic sources. The inconvenience, however, is that extracellular DNA, the most abundant component of soil DNA, is adsorbed by soil particles that are negatively charged and more resistant to DNase digestion than free DNA (Pietramellara et al., 2009; Taberlet et al., 2012). Finally on DNA degradation, Ushio et al., (2017) successfully recovered 200bp long fragments in pond water samples, but recovery of much larger fragments (i.e. 15kb) has been reported, also from water samples (Deiner, Renshaw, et al., 2017). In general, eDNA is detected for a longer period of time and tend to be of smaller size in soil than in water samples (Barnes & Turner, 2016).

In this study, we assessed the reliability of eDNA-based species detection for faunal diversity surveys. Using long-term camera-trapping data (Leempoel et al. 2019), we compared species identified from soil surface eDNA collected on trail segments located in front of 6 camera traps to species recorded by these cameras. We aimed to answer the following questions:

1. Can DNA from terrestrial mammals be recovered from soil surface samples and does it reliably reflect mammal diversity of that study area?

2. How do experimental parameters affect our ability to perform biodiversity monitoring using eDNA?

3. Are eDNA and camera trapping results comparable?

We collected large amounts of soil, extracted DNA and amplified target genes with mammal-specific primers. We then compared eDNA-detected species with the species diversity recorded by camera traps and other previous studies in the study area. As no experimental protocols have been defined so far, we compared different experimental parameters. In particular, we collected soil samples with two different sampling strategies, extracted DNA with two different extraction kits and performed PCR amplification with two metabarcodes of different sizes. We further evaluated quantitative relationships between eDNA and camera traps and looked at the temporal and spatial accuracy by comparing eDNA results with species distribution and abundance over three years.

## Methods

### Study area and camera trapping

Our study was conducted at the Jasper Ridge Biological Preserve (JRBP), California, USA, where we took advantage of a long-term camera trapping effort initiated in 2009. As of October 2017, the date of soil sampling, 18 wireless camera traps were installed, mostly along trails, to monitor wildlife (see Leempoel *et al.* 2019. for a detailed study based on these cameras). Out of these, we selected 6 cameras in contrasting habitats that recorded wildlife continuously for 1132 days before soil sampling (Fig S1).

Images from these cameras were manually examined by volunteers who identified animals and entered requisite metadata. We used these images for comparison with eDNA results. We first counted the number of cameras at which species were detected (i.e. occupancy) and calculated their relative abundance index (RAI) per day across the 6 cameras for periods of times ranging from 30 to 1132 days by steps of 60 days. We also gathered species presence/absence on a site-by-site basis for the same periods of time. Finally, we listed of all the mammals recorded by any of the cameras since 2009 for further comparison with eDNA data.

### Soil Sampling and DNA extraction

Two soil samples were collected at each site. For the first sample, we collected 20 cross-sections of the trail soil surface (first 2 cm), at 2m intervals centered in front of the camera, each filling a 50ml falcon tube, for a total of 1L mixed in a 1.6L sterile sampling bag (Fisher Scientific). For the second, we collected 80 subsamples in 12.5ml tubes every 2m for 80 meters in each direction from the camera, for a total of 1L. These subsamples consisted of random points either on the center or a side of the trail. Shovels were cleaned with bleach after each sample to prevent cross-contamination. Soil samples were frozen before extraction.

For DNA extraction, we used a dedicated pre-PCR laboratory room designed for low quality DNA samples that is separated from downstream PCR products. To avoid contamination, personnel were both physically and temporally separated from amplifications. We used two soil DNA extraction kits on the 12 soil samples collected. The first extraction protocol is based on the PowerMax (hereafter referred as PM) Soil DNA isolation Kit (MO BIO Laboratories, Inc.), as in Andersen *et al.* (2012). Mixed soils (10g) were processed following the manufacturer’s instructions. The second extraction protocol, developed by Taberlet *et al.* (2012) aimed to extract extracellular DNA by adding a saturated phosphate buffer (hereafter referred as PB) to the soil sample, followed by a filtration and elution using the NucleoSpin Soil kit (Macherey-Nagel, Düren, Germany), skipping the lysis step. We followed the protocol proposed by Taberlet *et al.* (2012), adding 1.97 g of NaH_2_PO_4_ and 14.7 g of Na_2_HPO_4_ to 1L of sterile water (Corning Cell Culture Grade Water, 25-055-CM) before mixing with 1L of soil sample in two sterile 1L bottles, and regularly shaking for 30 minutes. Two negative extractions were performed per extraction kit. The eluted DNA was quantified on a Nanodrop 2000 (Thermo Fisher Scientific Inc).

### DNA amplification and sequencing

We amplified mammal DNA with two metabarcodes of different size: a short ≈70bp metabarcode in the 16s (hereafter called M16s), and a longer ≈200bp metabarcode in the 12s (hereafter called M12s). The M16s was developed by Rasmussen *et al.* (2009) for human coprolite analysis and generally reaches its highest taxonomic resolution at the genus or family rank. This metabarcode was used by Andersen *et al.* (2012) to detect large mammals in both surface and core soil samples. The M12s has a higher resolution, reaching species rank for most sequences. It corresponds to the Mimammal-U developed by Ushio *et al.* (2017), who tested it on extracted DNA from 25 species representing major groups of mammals before testing it on pond water samples from zoo cages. For both metabarcodes, we used a 2-step PCR similar to Ushio *et al.* (2017). The primers are combined with 6 random bases and an adaptor in the 1^st^ PCR. Then the P5/P7 Miseq adaptors and dual-index barcodes (10 different forward and 13 reverse) are added to amplified sequences in a 2^nd^ PCR. For the 1^st^ PCR, we used 10μl of Amplitaq Gold 360 Master Mix, 1μl of each primer (5μM), 8μl mix template with H2O (PM: 8μl Template, PB: 4μl Template + 4μl H2O). Cycles: Holding 10min at 95°C, 45 cycles, denature 30s at 96°C, annealing for 30s at 60°C for M12s and 54°C for M16s, extension at 72°C for 60s for the M12s and 30s for the M16s, hold 10s at 72°C, and a final hold at 4°C. 3 μl of each PCR product were visualized on a 2% agarose gel and the remaining product was purified using the QIAquick PCR Purification Kit (Qiagen GmbH). For the 2^nd^ PCR, we used 10μl Amplitaq Gold 360 Master Mix, 1μl of each index primer, 3μl of template and 5μl H_2_O. Cycles: holding 10min at 95°C, 12 cycles, denature 30s at 96°C, annealing 30s at 65°C, extension 60s at 72°C, hold 10s at 72°C, hold at 4°C. Two negative PCRs were added for each primer pairs. The indexed second PCR products were quantified and assessed for quality control using an AATI Fragment Analyzer, normalized to equimolar concentrations and pooled together before purification using QIAquick PCR Purification Kit. Sequencing was performed in two separate runs with other unrelated projects on a MiSeq platform using the MiSeq Reagent Kit v3 (2 × 150-cycle) (Illumina, San Diego, CA, USA) with 30% PhiX and ran at the Stanford University PAN Facility.

### Sequence filtering and taxonomic assignment

We chose a series of filtering steps to be as conservative as possible while also attempting to retain as much of the ‘true’ eDNA diversity present in our soil samples. DNA sequences were automatically sorted (MiSeq post-processing) by amplicon pool using exact matches to the dual index barcodes. Then, sequences were filtered using the OBITools software (Boyer et al., 2016). Direct and reverse reads were aligned using *illuminapairedend*, and only sequences with a joined-alignment score above 40 were kept. Quality scores of paired sequences were checked using FastQC, prior to adapter trimming (a maximum mismatch of 10% with the primers was tolerated) in Cutadapt (Martin, 2011). At the same time, low quality sequences (quality score < 30) were removed. Afterwards, sequences shorter or longer than expected from the databases (see next paragraph) were removed using *Obigrep* (min. 24bp and max. 52bp for M16s, 150 and 192 for M12s). All samples were then pooled in a single fasta file and dereplicated using *Obiuniq*. Next, sequences occurring less than 10 times were removed before applying *Obiclean* to identify PCR and sequencing errors. To do so, *<u>Obiclean</u>* classifies sequences either as head, internal or singleton. Head sequences. the most common ones, correspond to true sequences or chimera product and can have multiple variants. Similarly, singletons are either true sequences or chimeras but are not related to any other sequences. Finally, internal sequences correspond to amplification/sequencing errors. See Boyer *et al.* (2016) for a more detailed explanation. *Obiclean* was applied sample by sample, with a maximum of one difference between two variant sequences and a threshold ratio between counts of one, meaning that all less abundant sequences are considered as variants. Only sequences with head or singleton status in at least one sample were kept. Further, sequences whose status in the global dataset was more commonly ‘internal’ than ‘head’ or ‘singleton’ were discarded (Giguet-Covex et al., 2014).

Afterwards, remaining sequences were matched against reference databases built using *EcoPCR* (Ficetola et al., 2010). To do so, we downloaded the EMBL database of standard sequences (http://ftp.ebi.ac.uk/pub/databases/embl/release/std/, release #135) of mammals (mam), vertebrates (vrt), mouse (mus) and human (hum), before converting it to the *EcoPCR* database format with *Obiconvert*. We then used *EcoPCR* to find sequences amplified by the primer pairs, using a maximum of 3 mismatches, and a minimum and maximum length identical to those mentioned above. Each resulting database was then dereplicated with *Obiuniq*. Expected mammal species were searched for in the database and their presence at species, genus or family rank was recorded to inform interpretation of results (See Supplementary Information). Thereafter, sequences were matched to the databases using *Ecotag*, and only sequences with an identity above 95% and 90% for the M12s and M16s respectively were kept. In addition, sequences that did not attain the rank of class or lower were deleted, and sequences assigned at species rank but whose identity was lower than 99% were ranked at genus level. Finally, sequences matching to the same reference sequence were grouped and their read count updated. After these steps, we proceeded on a case-by-case basis for regrouping or removal of sequences (Supplementary Information).

To discard potentially contaminant sequences, molecular operational taxonomic units (MOTUs) whose relative read abundance (RRA) was higher on average in the negative controls and/or negative PCRs than in true samples were removed. Finally, any MOTUs with abundances representing less than 0.05% of the total MOTU abundance across samples were removed to correct for potential cross-contamination. The detailed number of sequences, reads and taxa discarded at each step can be found in the Supplementary Information.

### Data analysis

We first compared the list of MOTUs reported by both metabarcodes and the number of samples in which they were detected. We calculated accumulation curves of the number of MOTUs as a function of the number of soil samples and PCR products with the function *specacccum* from the package vegan (Dixon, 2003) in R (R Development Core Team, 2018), using a 1000 permutations in the random method. Accumulation curves were used to compare between metabarcodes, sampling designs and extraction kits as a way to visualize putative total MOTU richness for each method.

We then compared the presence/absence of species from the eDNA survey with the camera trapping records. We performed linear regressions with the function “lm” in R between species occupancy and the RAI on the camera trap side, and the number of positive PCR, positive soil samples and positive sites on the eDNA side. Only species recorded by camera traps were kept for the comparison: black-tailed deer (*Odocoileus hemionus*), black-tailed jackrabbit (*Lepus* cf. *californicus*), bobcat (*Lynx rufus*), brush rabbit (*Sylvilagus bachmani*), coyote (*Canis* cf. *latrans*), mountain lion (*Puma concolor*), striped skunk (*Mephitis mephitis*), virginia opossum (*Didelphis* cf. *virginianus*). Note, however, that although we are confident with species-level identity in the camera images, not all these species can be identified down to the species level with either metabarcode, and this is denoted by our use of open nomenclature.

Finally, we calculated the similarity between eDNA and camera traps between matrices of species presence/absence per site (matrices of species X sites) from both camera traps and M12s eDNA using Mantel tests (Pearson method, 999 permutations) on dissimilarity indices calculated with vegdist, using the Jaccard method. With the same method, we calculated the similarity between the M12s and M16s by grouping MOTUs at the family level.

## Results

We obtained 9,795,610 and 6,008,091 paired reads for the M16s and M12s, respectively. After initial filtering, we identified 25 MOTUs with M16s and 44 MOTUs with M12s but many were removed or regrouped during our data processing pipeline. For example, all birds identified with M12s were removed as they were not part of the target taxon even though most of the bird species identified are indeed present in the community. In addition, multiple human haplotypes were detected and regrouped into a single MOTU. One MOTU was discarded based on a geographic criterion (*Capreolus* detected in M16s, a strictly Eurasian genus of deer, likely PCR error originating from Cervidae), and 3 had their taxonomy corrected because they are missing from the M16s database and were incorrectly assigned to a sister species or genus. We further detail the filtering procedure in the Supplementary Information. The proportion of human sequences was high, representing 49.14% in M16s and 35.30% in M12s. Humans are the most common species observed in the camera trap data, and although many sequences are likely to represent eDNA, there is also a high probability of human contamination in many phases of laboratory work. Thus, all human sequences were considered as contaminants in both metabarcodes. We also detected one additional contaminant, the gray fox (*Urocyon* cf. *cinereoargenteus*) in the M12s, for which we have no explanation except laboratory error. We note, however, that this contamination only occurred in one of the four negative extractions. After these final filtering steps, we detected 19 and 17 MOTUs with the short (M16s) and long (M12s) metabarcode respectively (Table 1).

**Table 1.**
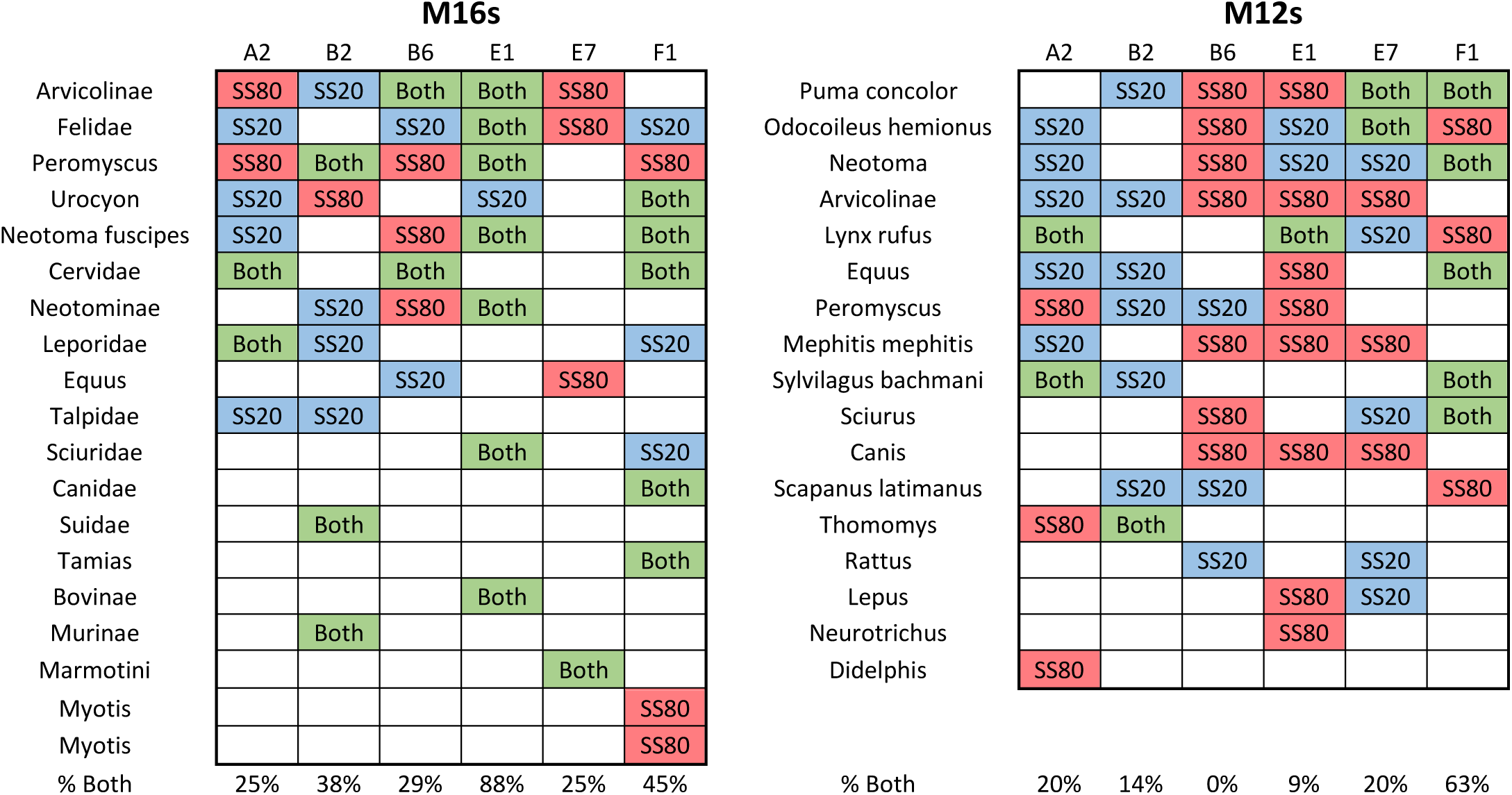
MOTUs identified by the short (M16s) and long (M12s) metabarcode at each site. “Both” indicates MOTUs detected in both soil samples collected at that site, whereas SS20 and SS80 indicates detection in the soil sample constituted of 20 or 80 subsamples respectively. % Both indicates how m any MOTUs at a given site were detected in both soil samples. MOTUs are ranked by the number of sites at which they were detected in decreasing order.

Species accumulation curves reached a maximum when accumulating the 36 PCR products per metabarcode, with a slightly faster progression for the M16s (Fig. 1a). By grouping PCRs per soil sample, however, curves are approaching a maximum with 12 soil samples and are similar for both metabarcodes (Fig 1b). Both metabarcodes also largely concurred at the Family level. Out of the 16 families detected, 10 were found with both metabarcodes and 3 were unique to each of them. The three families only detected using M16s were Bovinae, Suidae and Vespertilionidae (*Myotis* spp.), while for the M12s, these were *Mephitis mephitis* (Mephitidae), *Didelphis* cf. *virginianus*. (Didelphidae) and *Thomomys* cf. *bottae* (Geomyidae). However, the frequency at which MOTUs are detected by metabarcodes differed, as MOTUs detection from both soil samples of a given site were more frequent with the M16s than the M12s (43% and 20% respectively, Table 1). On the other hand, the maximum taxonomic resolution reached was higher for the M12s, with identification at genus rank or higher 16/17 times, compared to only 7/19 with the M16s. For example, *Puma concolor* and *Lynx rufus* identified at species level in M12s both correspond to Felidae in M16. There is a notable exception for the Sciuridae, for which we detected only one MOTU in M12s (*Sciurus* spp.) but three in M16s (*Tamias* spp., Sciuridae, Marmotini).

**Figure 1.**
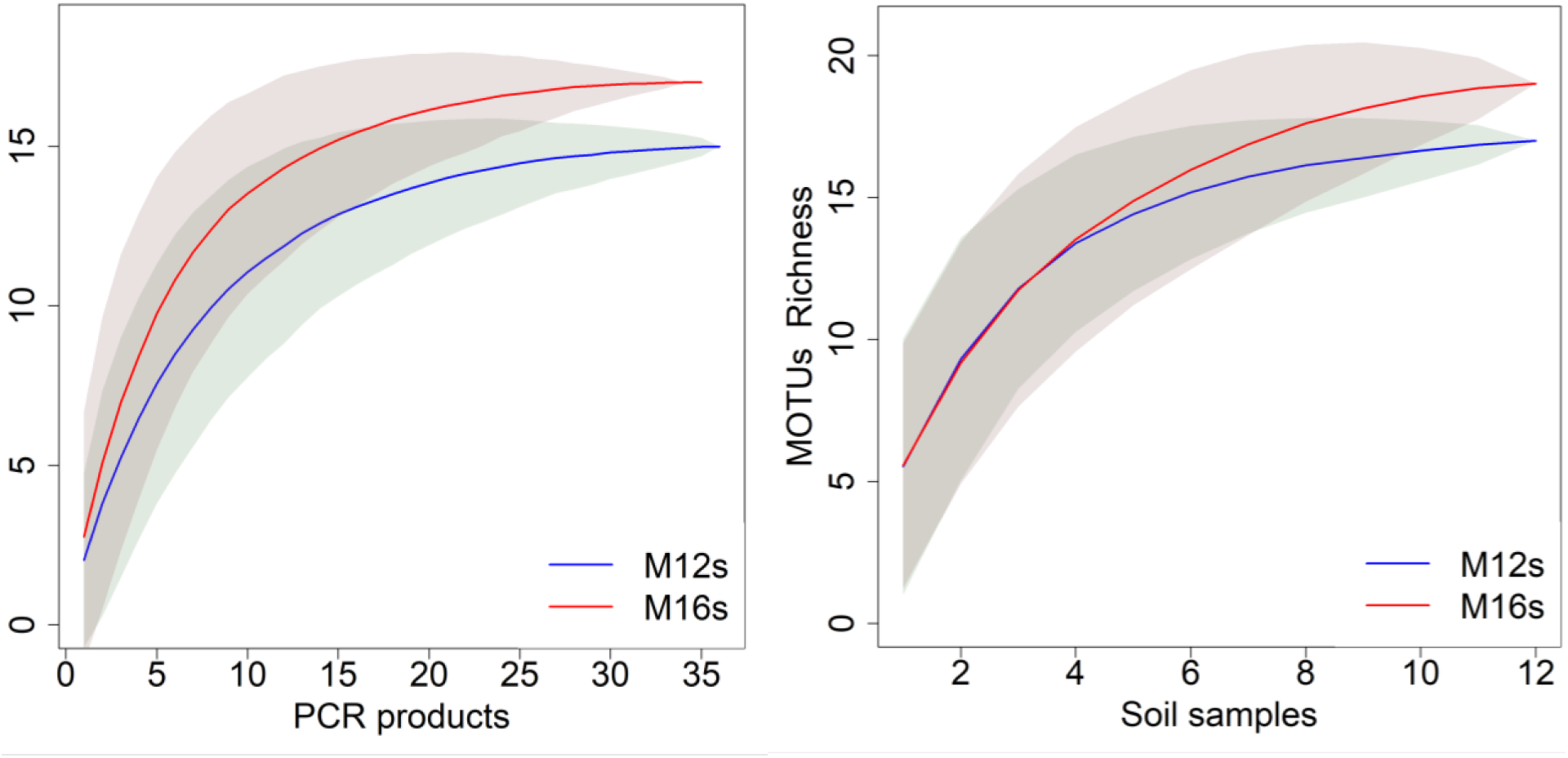
MOTUs accumulation curves per metabarcode. Based on MOTUs richness by PCR products (left) and soil sample (right, 2 samples of 1L collected per site).

There were no substantial differences between MOTUs obtained with the two different DNA extraction kits, as the majority of MOTUs were common in both (17/19 in M16s, 14/17 in M12s). In addition, MOTUs unique to one extraction protocol were rarely detected, between 1 and 3/36 PCR products (Marmotini and one of the two *Myotis* spp. only detected in PM extracts using M16s, and *Scapanus latimanus, Lepus* cf. *californicus* and *Didelphis* cf. *virginianus* only detected in PB extracts using M12s). For the 6 sites combined, soil samples made of 80 subsamples (SS80) slightly outperformed soil samples made of 20 subsamples (SS20) in the M12s but not in the M16s (Fig. S2). Here again, MOTUs unique to just one sampling design were among the rarest detected: *Myotis* spp. in M16s, *Neurotrichus* cf. *gibbsii* and *Didelphis* cf. *virginianus* in M12s are only found with SS80, while only *Rattus norvegicus* in M12s is found with SS20 only (Table 1).

All species recorded by camera traps were detected with eDNA, with the exception of the raccoon (*Procyon lotor*) (Fig 2). Conversely, all mid-to-large sized mammals detected with M12s were recorded by camera traps. Most importantly, a large panel of small mammals rarely, if ever, recorded by camera traps were detected with eDNA at multiple sites. All of them can be reasonably expected to be present in the preserve, as they have been documented by previous sightings and are known to inhabit the region (Jameson & Peeters, 2004; Young & Adams, 2009). With the M16s, however, two MOTUs not known to be in the preserve were detected (i.e. Suidae and Bovinae) in two soil samples but only from one site each.

**Figure 2.**
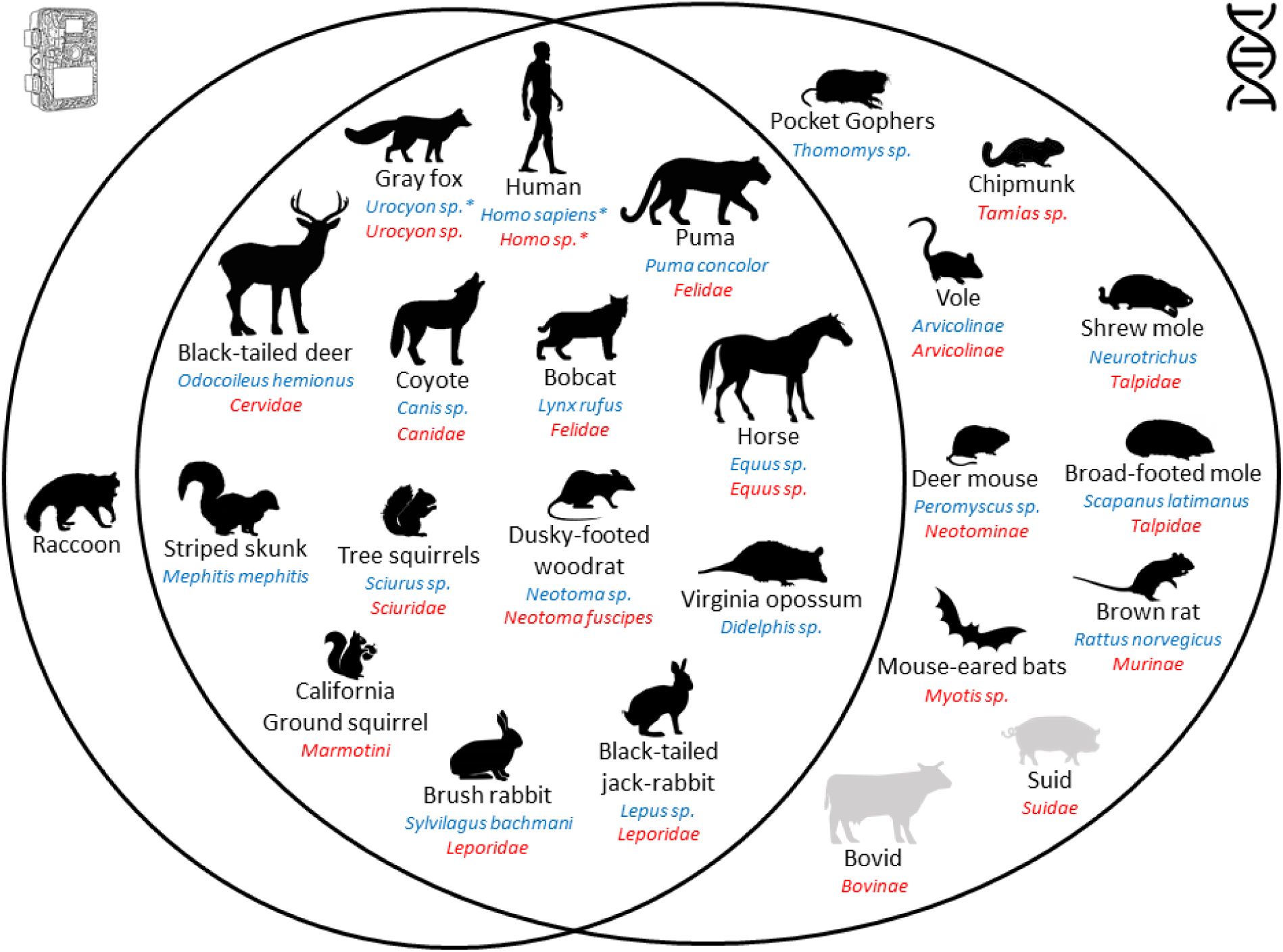
Venn Diagram between species recorded by camera traps and species detected with eDNA from both metabarcodes. Species detected with the M16s are marked in red and those detected with the M12s are marked blue. Scientific names are given at the maximum rank reached with eDNA. Species known to be present in the study area are in black. Species absent from the study area but detected with eDNA are in grey. Species considered as contaminant are indicated with *. Credits for illustration are in Supplementary Information.

We found a significant relationship between species occupancy from eDNA and camera traps. The number of sites at which species are recorded correlated with the number of sites at which they were detected with the M12s (Fig. 3). However, this relationship was only significant when using 30 days of camera-trapping data, and non-significant otherwise. All other attempts to identify a quantitative relationship between camera traps and eDNA were not significant, regardless of the length of camera-trapping survey chosen (Fig. S3-7).

**Figure 3.**
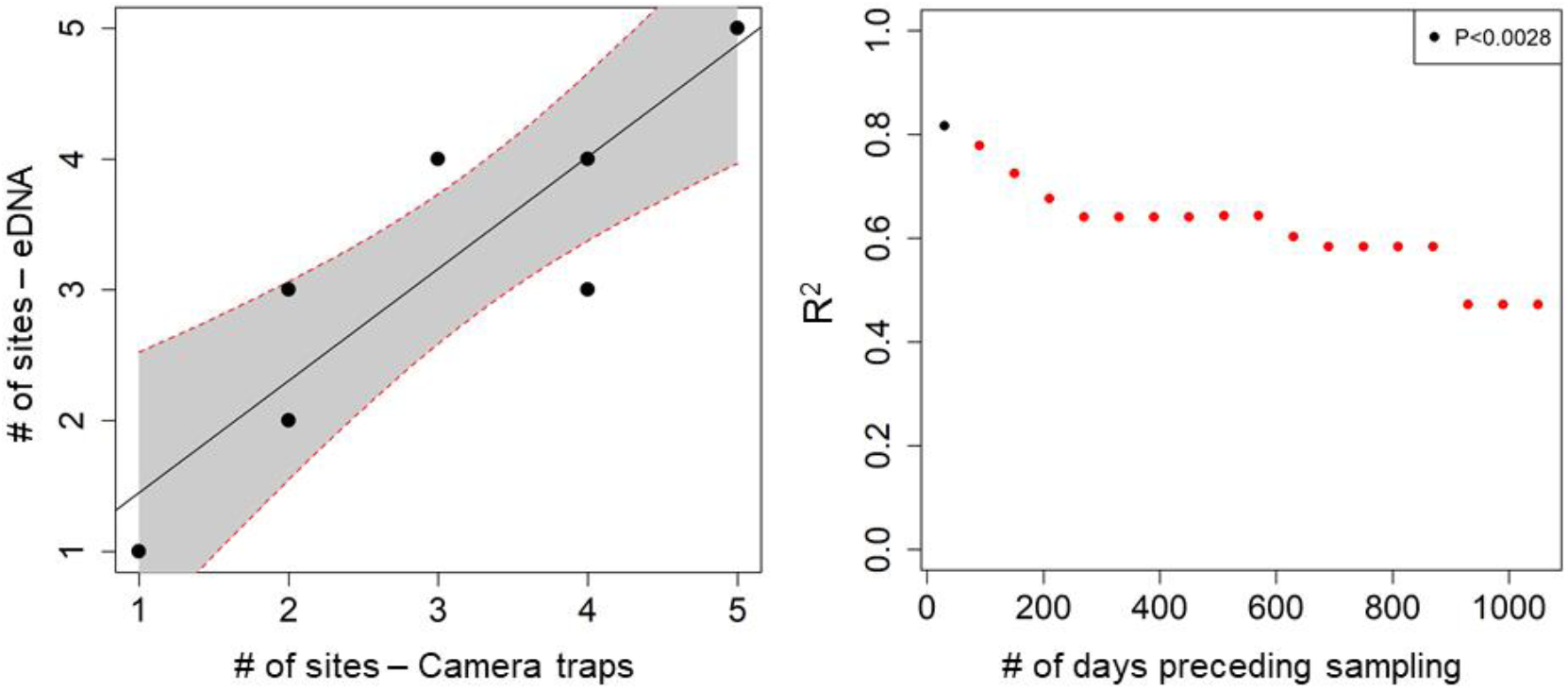
(Left) Quantitative relationship between the number of sites at which species (n= 9) are recorded by camera traps and detected with eDNA (M12s), using a month of camera trapping records preceding sampling. (Right) Change in the regression coefficient for the same relationship with increasing camera trap records. Camera trap records from 1 month to 3 years by steps of 60 days. Significant regressions (P<0.05 with Bonferroni correction) are marked with black points.

We also found substantial inconsistencies between the sites at which species are detected with eDNA and those at which they are recorded with camera traps. Indeed, the Mantel test between species presence/absence per site from camera trapping records against M12s detections was not significant (best correlation obtained with 180 days of camera trapping data, Mantel statistic: 0.485, significance: 0.097). Similarly, the Mantel test between family presence/absence per site from the M16s against M12s was not significant either (Mantel statistic: 0.487, Significance: 0.056).

## Discussion

We detected a large and similar ensemble of species with both metabarcodes, matching closely with the expected species composition in the studied area. Moreover, the detection of many small mammals is a considerable advantage of eDNA, as these species are generally more difficult to document than larger mammals. These results show that eDNA-based surveys could offer a meaningful alternative to the multiplicity of approaches that would have been needed to capture the same diversity.

We found several advantages to the longer metabarcode M12s (≈210bp) compared to the shorter M16s (≈70bp). First, the amount of long mitochondrial fragments we recovered from soil surface samples was sufficient to detect most species and attain high taxonomic rank in the majority of cases. Moreover, the sequence of the M12s can help us identify the most likely speices it belongs to. For example, multiple Arvicolinae are known in the region and the M16s sequence we collected does not go further than sub-family level and Blastn does not provide additional information. With the M12s Arvicolinae sequence, however, the top ten matches in Blastn are all from the genus microtus, suggesting the most likely candidate is *Microtus californicus*. Second, the M12s has the potential to discover previously undescribed haplotypes or subspecies. The *Neotoma* sequence we detected only matched at 98% with existing reference sequences of *Neotoma fuscipes*, and the only known occurrence of *Neotoma* in the region is the subspecies *Neotoma fuscipes annectens* (Matocq & Atocq, 2002). Third, the high taxonomic resolution of this metabarcode can help distinguish closely-related species. For example, the detection of *Rattus norvegicus* not only confirmed previous observations but also helps us differentiates it from another member of its genus, *Rattus rattus*, which is locally abundant in areas outside Jasper Ridge Biological Preserve but rare within it. Fourth, there were no unexpected taxa with the M12s unlike with the M16s. Bovinae and Suidae, detected only with the M16s, are not present in Jasper Ridge but can come from sources such as a: a nearby cattle ranch, contamination from food items or residuals of a large ranging predator diet. Wild boar, which is non-native but increasing its distribution in California was found in the puma diet in a parallel study at Jasper Ridge (In prep.). Anecdotally, we found that the M12s detected multiple species of birds, due to the lack of specificity of the Mimammal-U primers, suggesting that soil contains a large spectrum of above ground species that deserve further evaluation. While it could be seen as a disadvantage if one aims to maximize the number of reads for the target taxon, the amount of bird reads recovered was small (3%). The only major drawback to using such a large fragment is that we found them less frequently than short fragments, which could be an issue if the amount or quality of the starting material is limited.

Other experimental parameters studied had only a marginal influence on the final results. Regarding the use of two extraction kits, both performed equally well despite relying on very different protocols and amount of dirt processed. These results show that even for species with an expected low deposition rate, compared to plants or insects for example, the Phosphate Buffer (PB) extraction protocol is suitable. Moreover, PB costs a fraction of the price of the PowerMax (PM) extraction protocol and requires less equipment, making it ideal for extraction in the field (Taberlet et al., 2012; Zinger et al., 2016). In terms of sampling design, our results are in line with Andersen *et al.* (2012), who suggested collecting many small subsamples rather than a few large ones at each site. Our results also show that collecting a large amount of soil per site is important because of potentially high heterogeneity of soil samples and deposition rates (Taberlet et al., 2012; Yoccoz et al., 2012; Zinger et al., 2016). Here, 12L of surface soil was just enough to obtain a complete picture of the mammalian diversity. Finally, we sampled soil in all types of habitats in the preserve and noticed a higher number of detections at sites with shade and limited wind (Sites E1 and F1 in the riparian and oak-woodland habitats, Table 1), although we did not sample at enough sites per habitat to firmly support this observation.

Quantitative relationships of species abundance between eDNA and camera traps showed mixed results. We found a significant relationship between the number of sites where a species is seen by cameras and detected by eDNA, suggesting that eDNA could be used for species occupancy modelling, as is the case in cameras-trapping studies (O’Connell, Nichols, & Karanth, 2011). However, all other attempts failed. One of the key issues identified here is that, within a PCR product, 3 MOTUs usually represent more than 99% of the reads (Supplementary Information), suggesting that relative read abundance is an unlikely proxy for species abundance. Other authors have had better results with this metric in aquatic ecosystems (Hänfling et al., 2016; Thomsen et al., 2016), but there is a general lack of support in the literature for using relative read abundance in a quantitative way (Deiner, Bik, et al., 2017). Similarly, while we obtained an accurate picture of mammal diversity at the scale of the preserve, our eDNA results from a single site did not match camera trap images there. One reason for this mismatch could be that large parts of the trails covered during soil sampling are not covered by camera traps. However, the comparison between the M16s and M12s on a site by site analysis show a lack of similarity too. Future work should perhaps consider collecting larger volumes of soil per site but also process subsamples independently to find better correlations between species abundance and eDNA. Nevertheless, behavioral and ecological characteristics will also have a strong impact on the probability of detection with eDNA. In our data, some species appeared to be over-represented, such as the felines (puma, bobcat), which could be due to their habit of marking their territory via urine and feces along trails. Both felines are 3 and 10 times less frequent, respectively, on camera trap records than black-tailed deer whose feces are not typically found along trails, but were detected more frequently with eDNA.

Comparing with a long-term camera trapping study further allowed us to discuss the spatio-temporal accuracy of eDNA and the contribution of sighting frequency. We found that eDNA best reflected species presence for 30 to 150 days before sampling (Fig. 3b). In addition, coyotes were detected in only 3/12 soil samples, the least of all carnivores. Coyote, who were by far the dominant predator before the mountain lions increased substantially in 2013 (Leempoel et al., 2019), but now are seen an order of magnitude less frequently (984 records in 2012, 98 in 2017). This suggest that the DNA of coyotes does not stay detectable for long (4+ years) in the environment and that recent abundance does affect the probability of detection in eDNA. Similarly, the relative abundance of racoons decreased substantially over the past several years (150 records in 2012 to 24 in 2017) and we were not able to detect them at any site using our eDNA methods. We also did not detect the American badger (*Taxidea taxus*) nor the long-tailed weasel (*Mustela frenata*), but we have recorded just 3 pictures for each of them since 2009, and these images were not taken from any of the 6 sites sampled. Overall, our results suggest that the DNA of terrestrial mammals on the soil surface does not last more than a couple of years, remains local, and is not generally detectable for extremely rare species using our protocols.

While these results are promising for eDNA as a survey tool for terrestrial mammals, such studies need to be improved and replicated in many habitats and environments before being considered for ecosystem surveys more globally. The major and critical hindrances we faced were the incompleteness of reference databases and improper amplification due to primer mismatch. At least two inconsistent results between both metabarcodes can be attributed to these factors. For example, *Didelphis* and *Thomomys* are not present in the reference database for the M16s. For *Didelphis*, this is due to too high a mismatch with the M16s forward primer, while for *Thomomys*, it is due to the absence of a reference sequence for any sister species in its family (Geomyidae). If this data shortfall is an issue in a region where biodiversity is well studied, it is easy to imagine that it can only be worse in poorly covered ecosystems and/or more biodiverse regions. To investigate the utility of our approach more globally and using the same methods as described in Supplementary Information, we found that 59% of all known mammals (IUCN, 2019) are missing from the M12s database (Table S1). Indeed, 33% of these missing species have no sister species at the genus level. These numbers get even worse for specific orders, with 48% of missing rodents having no sister species at the genus level or 62% for missing carnivores. This can only be more problematic for other groups of animals and it greatly hampers our ability to conduct eDNA studies worldwide. Moreover, this preliminary analysis of detectability and database evaluation is often overlooked in eDNA papers, despite being a critical step to understand if the non-detection of species is due to their true absence in the studied area or to the shortcomings of databases.

Another consequence of incomplete databases is that sequences with an identity below the defined filtering threshold will be discarded. We therefore looked at these discarded sequences to find potentially missing species. We found a sequence corresponding to the sub-family Soricinae in the M16s sequences, discarded because its best identity was lower that the decided threshold (i.e. 90%). This sequence likely corresponds to *Sorex ornatus*, the only known *Sorex* in the preserve. Similarly, several sequences attributed to squirrels in the M12s were discarded, which could help explain why we did not successfully report their known diversity in the study area. For example, we found a sequence attributed to *Callospermophilus lateralis*, but instead it likely belongs to the California ground squirrel (*Otospermophilus beecheyi*), which is abundant in the preserve, but also is missing from the M12s database at the species and genus level. We also note that the only sequence kept and attributed to squirrels in the M12s results had a maximum identity of 96% with *Scurius niger* using BLASTn, suggesting the genus is correct but not the species. Two other *Sciurus* are known in the area, *Sciurus griseus* and *Sciurus carolinensis*, but they are not in the reference database either. It is worth mentioning that 2 MOTUs of bats were detected out of the 14 known in the region, showing the limit of eDNA for this order of mammals.

Part of the success of this study can be attributed to the habitat and topography of Jasper Ridge. Indeed, the vegetation is a dense chaparral in many areas, discouraging the movement of large mammals. Steep, uneven terrain, deep drainages, and creeks that flood in the rainy season also influence the routes animals take across the landscape. Therefore, trails represent the easiest way to move around the reserve. Thus, large mammals repeatedly pass by the same locations, increasing the concentration of DNA dropped on trails. Most of our cameras are set on trails or at trail intersections for this reason. This opens the question of whether we would have been able to detect these species with eDNA had we sampled randomly in an open grassland with no clear trail structure. Still, small mammals, who do not rely as much on trails, were easily detected with eDNA.

Our study demonstrates once more that eDNA is a remarkably promising approach for ecosystem surveys and opens new possibilities for managers and researchers to reveal the distribution and interaction of species in a single survey. We detected the vast majority of species present in the study area, including those which are generally too small to trigger camera traps. As such, eDNA alone was sufficient to obtain a reasonable picture of species diversity, outperformed camera or live trapping, and did not require previous knowledge of the study area. We further suggest that long fragments (200bp) are ideal for present-day biodiversity studies, as they are less likely than short fragments (70bp) to last for years in the top soil layer yet frequently recovered. However, as long as mitochondrial reference databases remain so incomplete, eDNA can only be considered as a complement to existing approaches rather than an alternative.

## Acknowledgments

Swiss National Science Foundation Early Postdoc Mobility grant P2ELP3_175075, Stanford University Mellon grant (KL), HHMI Professorship (EAH). We thank many Jasper Ridge volunteers for their contribution to the tagging of camera trap images, Hadly lab members for their suggestions, Fred Boyer for helping us troubleshoot with OBITools, Nicole Nova for providing several illustrations of mammals.

## Author contribution

K.L. designed and performed the research, analyzed the data and wrote the paper. T.H. installed and maintained the camera traps, curated the image database and contributed to the paper. E.A.H designed the research and contributed to the paper.

## References

Adams, I. C., Hoekstra, A. L., Muell, R. M., & Janzen, J. F. (2019). A Brief Review of Non-Avian Reptile Environmental DNA (eDNA), with a Case Study of Painted Turtle (Chrysemys picta) eDNA Under Field Conditions. Diversity, Vol. 11. doi: 10.3390/d11040050

Andersen, K., Bird, K. L., Rasmussen, M., Haile, J., Breuning-Madsen, H., Kjær, K. H., … Willerslev, E. (2012). Meta-barcoding of “dirt” DNA from soil reflects vertebrate biodiversity. Molecular Ecology, 21(8), 1966–1979. doi: 10.1111/j.1365-294X.2011.05261.x

Barnes, M. A., & Turner, C. R. (2016). The ecology of environmental DNA and implications for conservation genetics. Conservation Genetics, Vol. 17, pp. 1–17. doi: 10.1007/s10592-015-0775-4

Barnosky, A. D., Hadly, E. a, Bascompte, J., Berlow, E. L., Brown, J. H., Fortelius, M., … Smith, A. B. (2012). Approaching a state shift in Earth’s biosphere. Nature, 486(7401), 52–58. doi: 10.1038/nature11018

Bickford, D., Lohman, D. J., Sodhi, N. S., Ng, P. K. L., Meier, R., Winker, K., … Das, I. (2007). Cryptic species as a window on diversity and conservation. Trends in Ecology and Evolution, Vol. 22, pp. 148–155. doi: 10.1016/j.tree.2006.11.004

Bovendorp, R. S., McCleery, R. A., & Galetti, M. (2017). Optimising sampling methods for small mammal communities in Neotropical rainforests. Mammal Review. doi: 10.1111/mam.12088

Boyer, F., Mercier, C., Bonin, A., Le Bras, Y., Taberlet, P., & Coissac, E. (2016). obitools: A unixinspired software package for DNA metabarcoding. Molecular Ecology Resources. doi: 10.1111/1755-0998.12428

Burton, A. C., Neilson, E., Moreira, D., Ladle, A., Steenweg, R., Fisher, J. T., … Boutin, S. (2015). Wildlife camera trapping: A review and recommendations for linking surveys to ecological processes. Journal of Applied Ecology, Vol. 52, pp. 675–685. doi: 10.1111/1365-2664.12432

Deiner, K., Bik, H. M., Mächler, E., Seymour, M., Lacoursière-Roussel, A., Altermatt, F., … Bernatchez, L. (2017). Environmental DNA metabarcoding: Transforming how we survey animal and plant communities. Molecular Ecology. doi: 10.1111/mec.14350

Deiner, K., Renshaw, M. A., Li, Y., Olds, B. P., Lodge, D. M., & Pfrender, M. E. (2017). Long-range PCR allows sequencing of mitochondrial genomes from environmental DNA. Methods in Ecology and Evolution. doi: 10.1111/2041-210X.12836

Dixon, P. (2003). VEGAN, a package of R functions for community ecology. Journal of Vegetation Science, 14, 927–930. doi: 10.1111/j.1654-1103.2003.tb02228.x

Drummond, A. J., Newcomb, R. D., Buckley, T. R., Xie, D., Dopheide, A., Potter, B. C. M., … Nelson, N. (2015). Evaluating a multigene environmental DNA approach for biodiversity assessment. GigaScience. doi: 10.1186/s13742-015-0086-1

Ficetola, G. F., Coissac, E., Zundel, S., Riaz, T., Shehzad, W., Bessière, J., … Pompanon, F. (2010). An In silico approach for the evaluation of DNA barcodes. BMC Genomics. doi: 10.1186/1471-2164-11-434

Frishkoff, L. O., Karp, D. S., M’Gonigle, L. K., Mendenhall, C. D., Zook, J., Kremen, C., … Daily, G. C. (2014). Loss of avian phylogenetic diversity in neotropical agricultural systems. Science, 345(6202), 1343–1346. doi: 10.7910/DVN/26910

Giguet-Covex, C., Pansu, J., Arnaud, F., Rey, P. J., Griggo, C., Gielly, L., … Taberlet, P. (2014). Long livestock farming history and human landscape shaping revealed by lake sediment DNA. Nature Communications. doi: 10.1038/ncomms4211

Hänfling, B., Handley, L. L., Read, D. S., Hahn, C., Li, J., Nichols, P., … Winfield, I. J. (2016). Environmental DNA metabarcoding of lake fish communities reflects long-term data from established survey methods. Molecular Ecology. doi: 10.1111/mec.13660

Harkins, K. M., Keinath, D., & Ben-David, M. (2019). It’s a trap: Optimizing detection of rare small mammals. PLOS ONE, 14(3), e0213201. Retrieved from https://doi.org/10.1371/journal.pone.0213201

Ishige, T., Miya, M., Ushio, M., Sado, T., Ushioda, M., Maebashi, K., … Matsubayashi, H. (2017). Tropical-forest mammals as detected by environmental DNA at natural saltlicks in Borneo. Biological Conservation, 210, 281–285. doi: 10.1016/j.biocon.2017.04.023

IUCN. (2019). The IUCN Red List of Threatened Species. Retrieved May 5, 2019, from http://www.iucnredlist.org

Jameson, E. W., & Peeters, H. J. (2004). Mammals of California. Berkeley: University of California Press.

Leempoel, K., Meyer, J. M., Hebert, T., Nova, N., & Hadly, E. A. (2019). Return of an apex predator to a suburban preserve triggers a rapid trophic cascade. BioRxiv, 564294. doi: 10.1101/564294

Martin, M. (2011). Cutadapt removes adapter sequences from high-throughput sequencing reads. EMBnet.Journal. doi: 10.14806/ej.17.1.200

Matocq, M. D., & Atocq, M. A. D. M. (2002). Morphological and molecular analysis of a contact zone in the Neotoma fuscipes species complex. Journal of Mammalogy.

Mendenhall, C. D., Karp, D. S., Meyer, C. F. J., Hadly, E. A., & Daily, G. C. (2014). Predicting biodiversity change and averting collapse in agricultural landscapes. Nature, 509(7499), 213–217. doi: 10.1038/nature13139

Mendenhall, C. D., Shields-Estrada, A., Krishnaswami, A. J., & Daily, G. C. (2016). Quantifying and sustaining biodiversity in tropical agricultural landscapes. Proceedings of the National Academy of Sciences of the United States of America, 201604981. doi: 10.1073/pnas.1604981113

O’Connell, A. F., Nichols, J. D., & Karanth, K. U. (2010). Camera traps in animal ecology: methods and analyses. Springer Science & Business Media.

O’Connell, A. F., Nichols, J. D., & Karanth, K. U. (2011). Camera traps in animal ecology: Methods and analyses. In Camera Traps in Animal Ecology: Methods and Analyses. doi: 10.1007/978-4-431-99495-4

Pietramellara, G., Ascher, J., Borgogni, F., Ceccherini, M. T., Guerri, G., & Nannipieri, P. (2009). Extracellular DNA in soil and sediment: Fate and ecological relevance. Biology and Fertility of Soils, Vol. 45, pp. 219–235. doi: 10.1007/s00374-008-0345-8

Port, J. A., O’Donnell, J. L., Romero-Maraccini, O. C., Leary, P. R., Litvin, S. Y., Nickols, K. J., … Kelly, R. P. (2016). Assessing vertebrate biodiversity in a kelp forest ecosystem using environmental DNA. Molecular Ecology, 25(2), 527–541. doi: 10.1111/mec.13481

R Development Core Team. (2018). R: A Language and Environment for Statistical Computing.

Rasmussen, M., Cummings, L. S., Gilbert, M. T. P., Bryant, V., Smith, C., Jenkins, D. L., & Willerslev, E. (2009). Response to comment by Goldberg et al. on “DNA from pre-clovis human coprolites in Oregon, North America.” Science. doi: 10.1126/science.1167672

Stem, C., Margoluis, R., Salafsky, N., & Brown, M. (2005). Monitoring and evaluation in conservation: A review of trends and approaches. Conservation Biology, Vol. 19, pp. 295–309. doi: 10.1111/j.1523-1739.2005.00594.x

Tabak, M. A., Norouzzadeh, M. S., Wolfson, D. W., Sweeney, S. J., Vercauteren, K. C., Snow, N. P., … Miller, R. S. (2018). Machine learning to classify animal species in camera trap images: Applications in ecology. Methods in Ecology and Evolution. doi: 10.1111/2041-210X.13120

Taberlet, P., Prud’Homme, S. M., Campione, E., Roy, J., Miquel, C., Shehzad, W., … Coissac, E. (2012). Soil sampling and isolation of extracellular DNA from large amount of starting material suitable for metabarcoding studies. Molecular Ecology, 21(8), 1816–1820. doi: 10.1111/j.1365-294X.2011.05317.x

Thomsen, P. F., Kielgast, J., Iversen, L. L., Moller, P. R., Rasmussen, M., & Willerslev, E. (2012). Detection of a Diverse Marine Fish Fauna Using Environmental DNA from Seawater Samples. PLoS ONE, 7(8). doi: 10.1371/journal.pone.0041732

Thomsen, P. F., Møller, P. R., Sigsgaard, E. E., Knudsen, S. W., Jørgensen, O. A., & Willerslev, E. (2016). Environmental DNA from seawater samples correlate with trawl catches of subarctic, deepwater fishes. PLoS ONE. doi: 10.1371/journal.pone.0165252

Ushio, M., Fukuda, H., Inoue, T., Makoto, K., Kishida, O., Sato, K., … Miya, M. (2017). Environmental DNA enables detection of terrestrial mammals from forest pond water. Molecular Ecology Resources. doi: 10.1111/1755-0998.12690

Valentini, A., Taberlet, P., Miaud, C., Civade, R., Herder, J., Thomsen, P. F., … Dejean, T. (2016). Next-generation monitoring of aquatic biodiversity using environmental DNA metabarcoding. Molecular Ecology, 25(4), 929–942. doi: 10.1111/mec.13428

Yoccoz, N. G., Brathen, K. A., Gielly, L., Haile, J., Edwards, M. E., Goslar, T., … Taberlet, P. (2012). DNA from soil mirrors plant taxonomic and growth form diversity. Molecular Ecology, 21(15), 3647–3655. doi: 10.1111/j.1365-294X.2012.05545.x

Young, H., & Adams, R. (2009). Small Mammals at Jasper Ridge.

Zinger, L., Chave, J., Coissac, E., Iribar, A., Louisanna, E., Manzi, S., … Taberlet, P. (2016). Extracellular DNA extraction is a fast, cheap and reliable alternative for multi-taxa surveys based on soil DNA. Soil Biology and Biochemistry. doi: 10.1016/j.soilbio.2016.01.008

