## Supplementary information for "A comparison of eDNA to camera trapping for assessment of terrestrial mammal diversity"

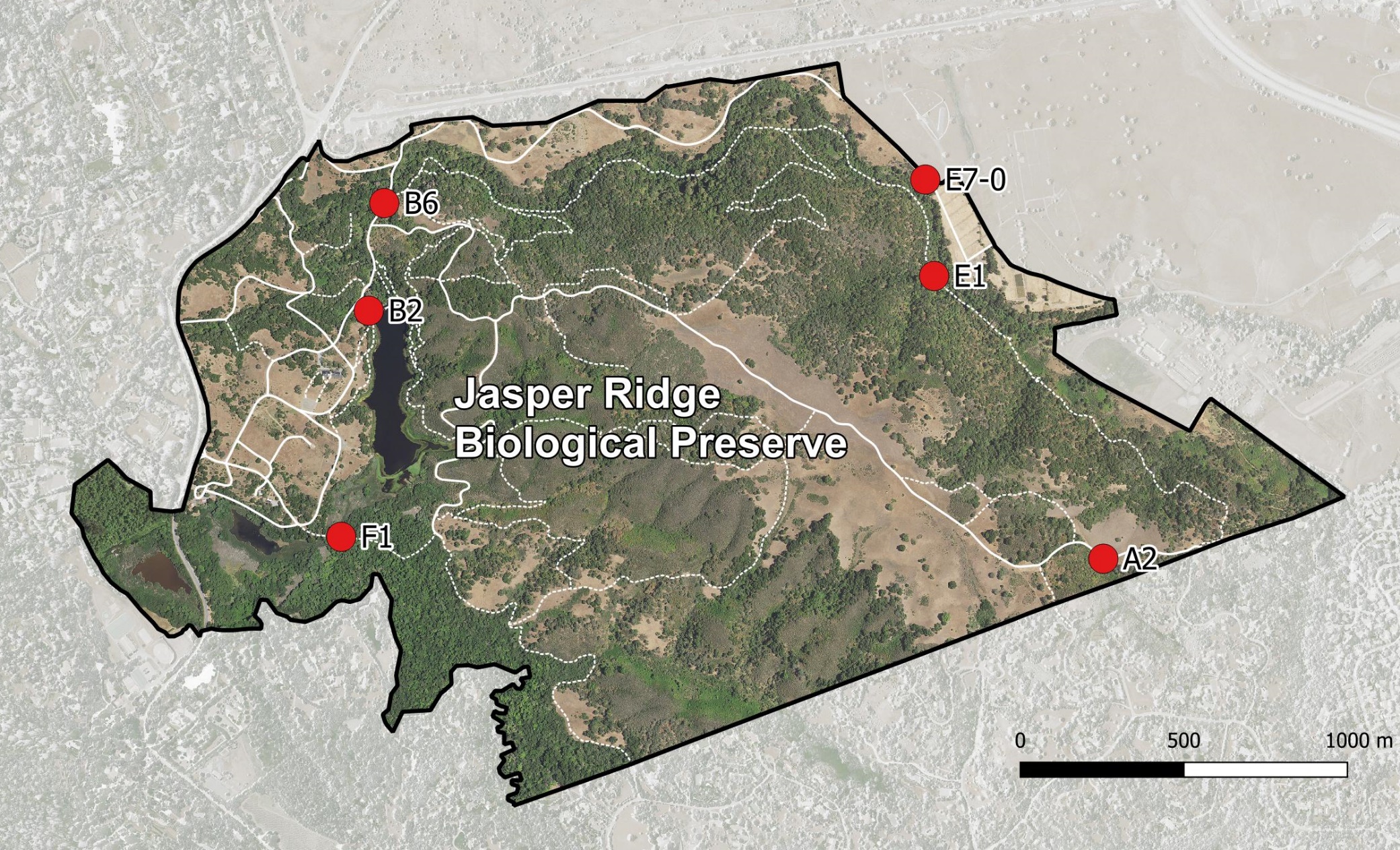

Figure S1 Map of Jasper Ridge Biological Preserve and the 7 sampling sites corresponding to camera traps (6 soil sampling sites and the spring).

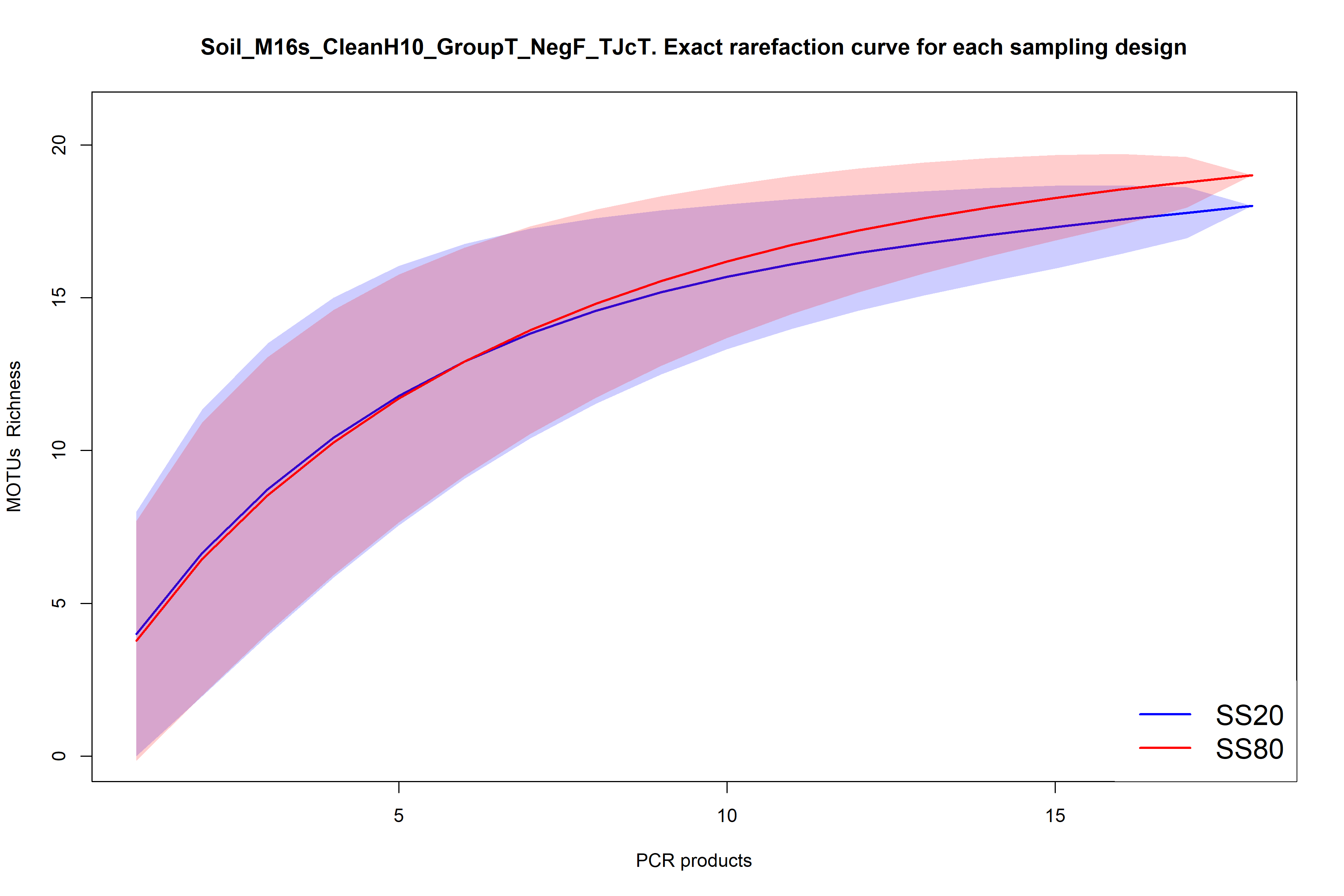

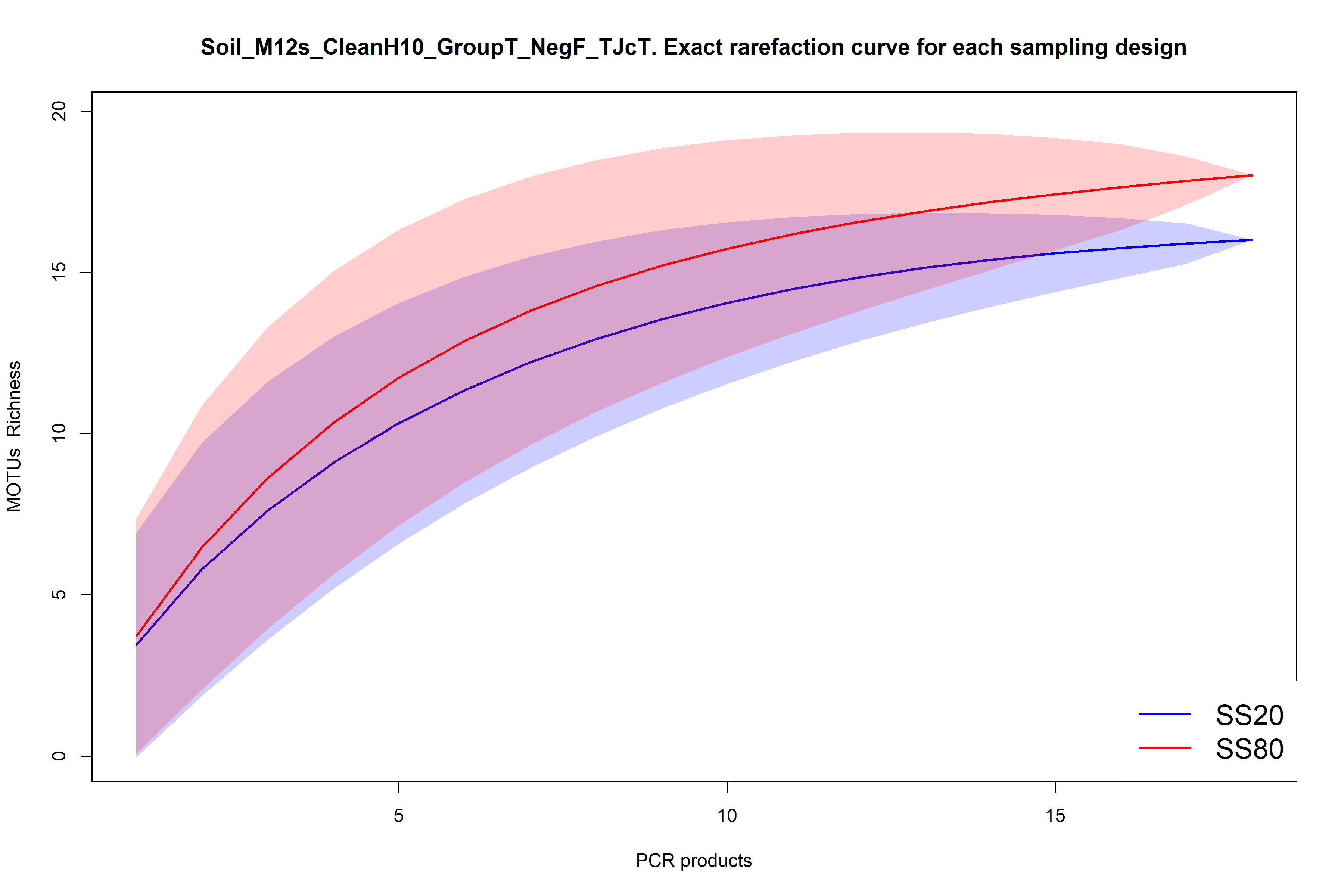

Figure S2 MOTUs accumulation curves per soil sampling strategy for the M16s (top) and M12s (bottom).

**MOTUs removed from the M12s results (M12s Filtered Ecotag spreadsheet)**

1. Removing birds: GQ264618, AY164517, AF407087, EF153719, KR732817, JX516068, KM078763, AF447214
2. Grouping scurius niger sequences: U59174, U67289
3. Removing hominidae sequences not as species level: AC008434,AC197159,D38116
4. Grouping homo sapiens sequences: AB196722, AC002087, AC021914, AC062033, AC074322, AC090204
5. Likely PCR errors in Cricetidae removed: JF261174, KY707300, KY707303, KY707310
6. Grouping horse sequences: KT368731,AP013078
7. Likely PCR error. Removing Felinae sequences that should have reached higher taxonomic level given that all known Felinae in the region are sequenced: AY012150, KM224285.
8. Unclear origin, removed. Only matches to 1 sequence of canis lupus familiaris on Blastn but it is not clear for which gene: AB048589
9. Likely PCR error. Removing Odocoileus sequences that should have reached higher taxonomic level given that all known Odocoileus in the region are sequenced: JN632672

**MOTUs removed from the M16s results (M16s Filtered Ecotag spreadsheet)**

1. Grouping Equus sequences: HQ439494, AB859014
2. Taxonomic correction. Changing taxa name for Peromyscus since this species is not in the region. Likely that Peromyscus in the region have the same sequence: KY707308
3. Likely PCR error removed. Capreolus, genus of Eurasian deer: AY122048
4. Taxonomic correction. Genus not in the region but its tribe is the same as the California ground squirrel: KP698974
5. Taxonomic correction. Genus Scalopus not in the region but there are Talpidae in Jasper Ridge: AF069539
6. Sequence removed, likely PCR error. Blastn of the sequence shows it also matches 100% with known human sequences: AC146433
7. Grouping Homo sequences: AB017708, AC021451, AC024248

**Credits of illustrations from Figure 2**

Striped skunk: Kevin Leempoel

Black-tailed deer: Nicole Nova

Mountain lion: Nicole Nova

Coyote: Nicole Nova

Bobcat: Nicole Nova

Gray fox: Nicole Nova

[Raccoon: http://phylopic.org/image/52193065-f85e-4e4d-a677-11d08aed8c2e/](Raccoon:%20http://phylopic.org/image/52193065-f85e-4e4d-a677-11d08aed8c2e/)

Horse: <http://phylopic.org/image/d0847124-56fe-4ca5-8415-e75b0afdc751/>

Black-tailed jackrabbit: <http://phylopic.org/image/8e61e166-11f4-4377-a923-9b5b597b6eba/>

Brush rabbit: <http://phylopic.org/image/dea688b6-9168-4e79-a106-366888148eb1/>

Squirrel: <http://phylopic.org/image/dad08fea-5263-4f57-a37b-c27cbe0eb9a5/>

Opossum: Sarah Werning; <http://creativecommons.org/licenses/by/3.0/>; <http://phylopic.org/image/91324e57-b3f1-42e0-abe3-43e5bc8aa4c6/>

Pocket gopher: Kevin Leempoel

Chipmunk: [Freepik] from [www.flaticon.com](http://www.flaticon.com)

Vole: <http://phylopic.org/image/ba1d31e7-2679-418b-9df6-71dbbbac78d9/>

Deer mouse: <http://phylopic.org/image/81930c02-5f26-43f7-9c19-e9831e780e53/>

Shrew Mole: [Freepik] <https://www.flaticon.com/free-icon/mole_371930#term=mole&page=1&position=37>

Broad-footed mole: Kevin Leempoel

Mouse-eared bats: [Freepik] <https://www.flaticon.com/free-icon/flying-bat_12311#term=bat&page=1&position=71>

Norwegian rat: Rebecca Groom, <http://creativecommons.org/licenses/by-sa/3.0/>, <http://phylopic.org/image/726683a1-78ed-4371-9c10-c220b8716c54/>

Bovine: [Freepik] <https://www.flaticon.com/free-icon/cow-silhouette_62470#term=cow&page=1&position=1>

Suidae: [Freepik] <https://www.flaticon.com/free-icon/pig-side-view-silhouette_47071#term=pig&page=1&position=24>

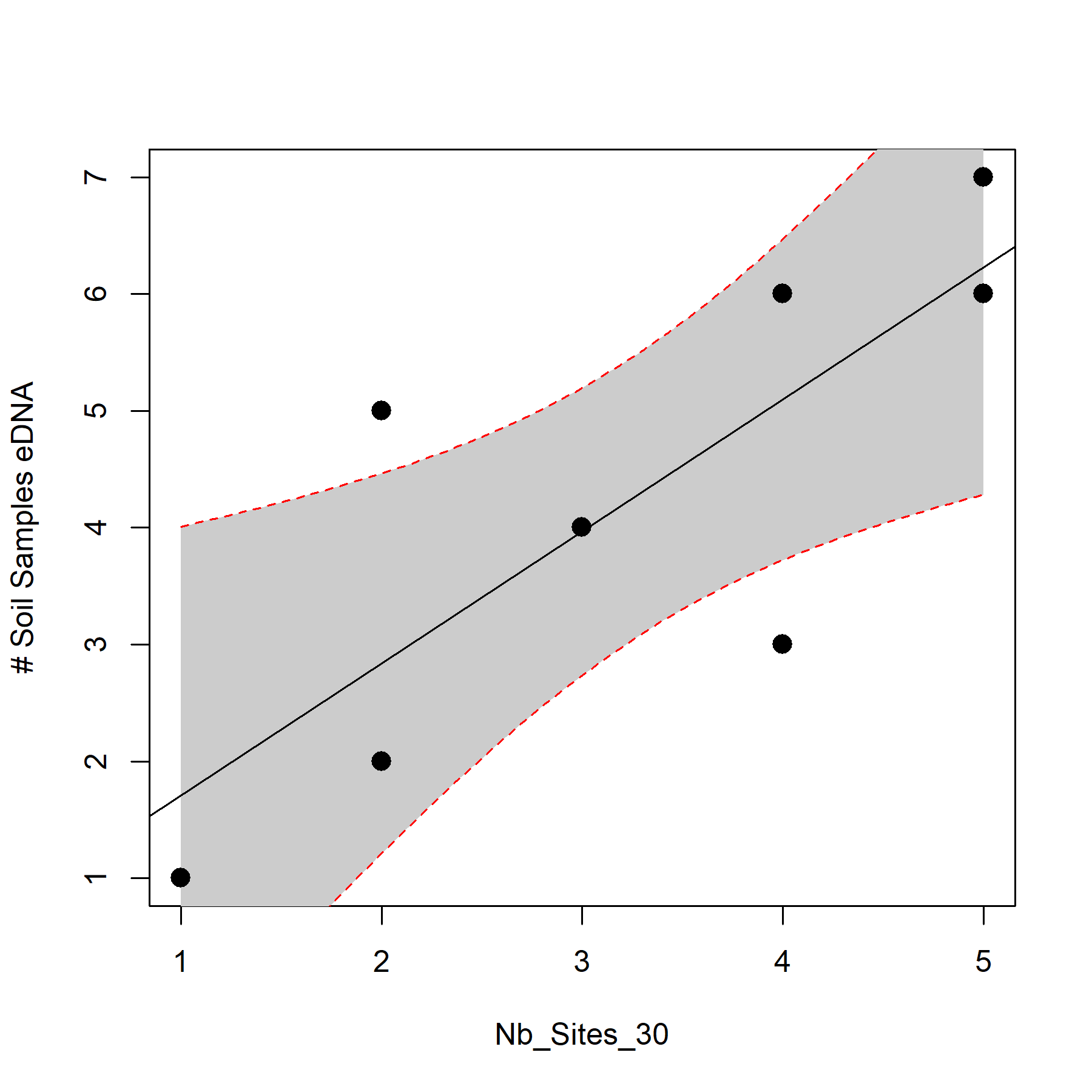

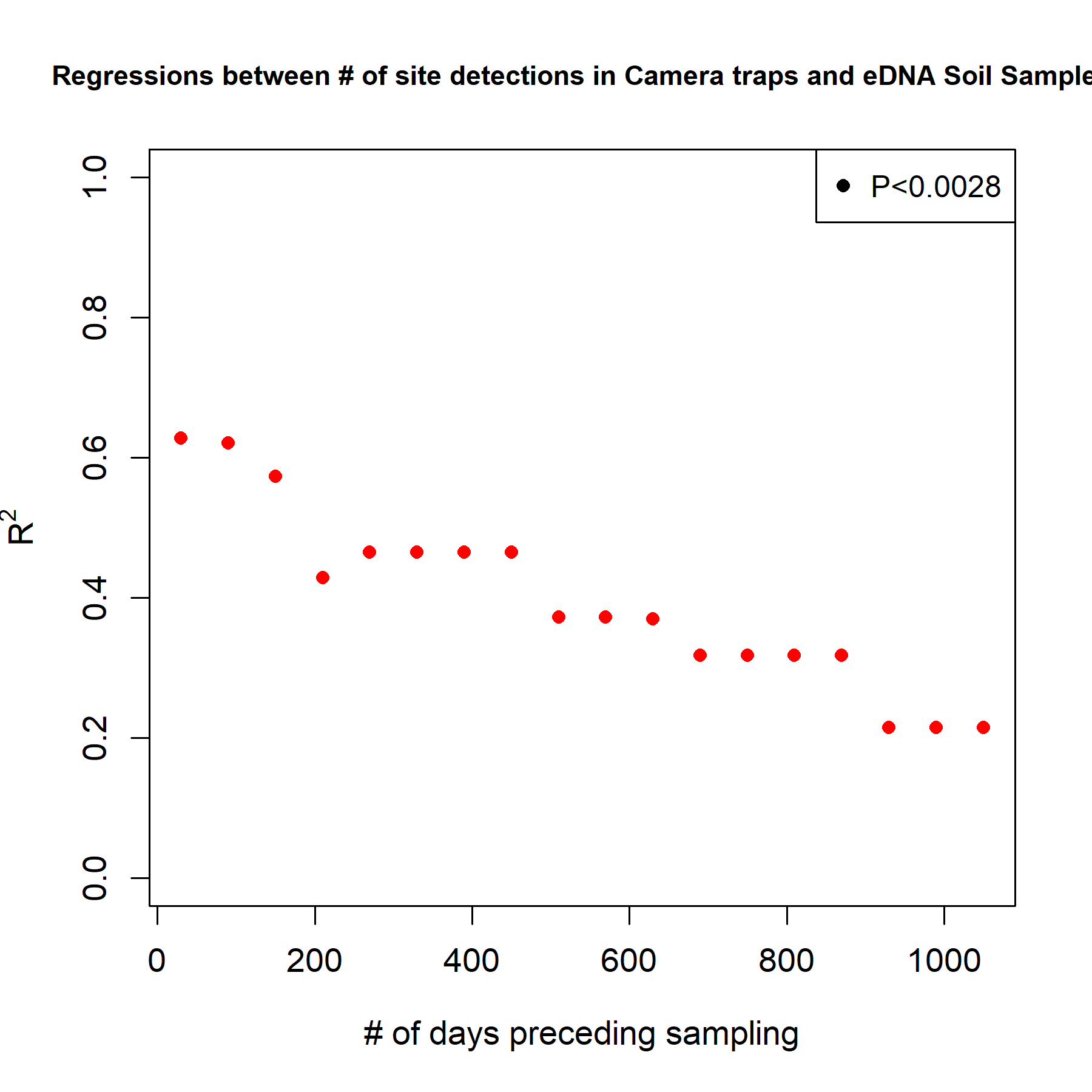

Figure S3 (Top) Quantitative relationship between the number of sites at which species (n=9) are recorded by camera traps and detected in eDNA soil samples (M12s), using a month of camera trapping records preceding sampling. (Bottom) Change in the regression coefficient for the same relationship with increasing camera trap records. Camera trap records from 1 month to 3 years by steps of 60 days. Significant regressions (P<0.05 with Bonferroni correction) are marked with black points.

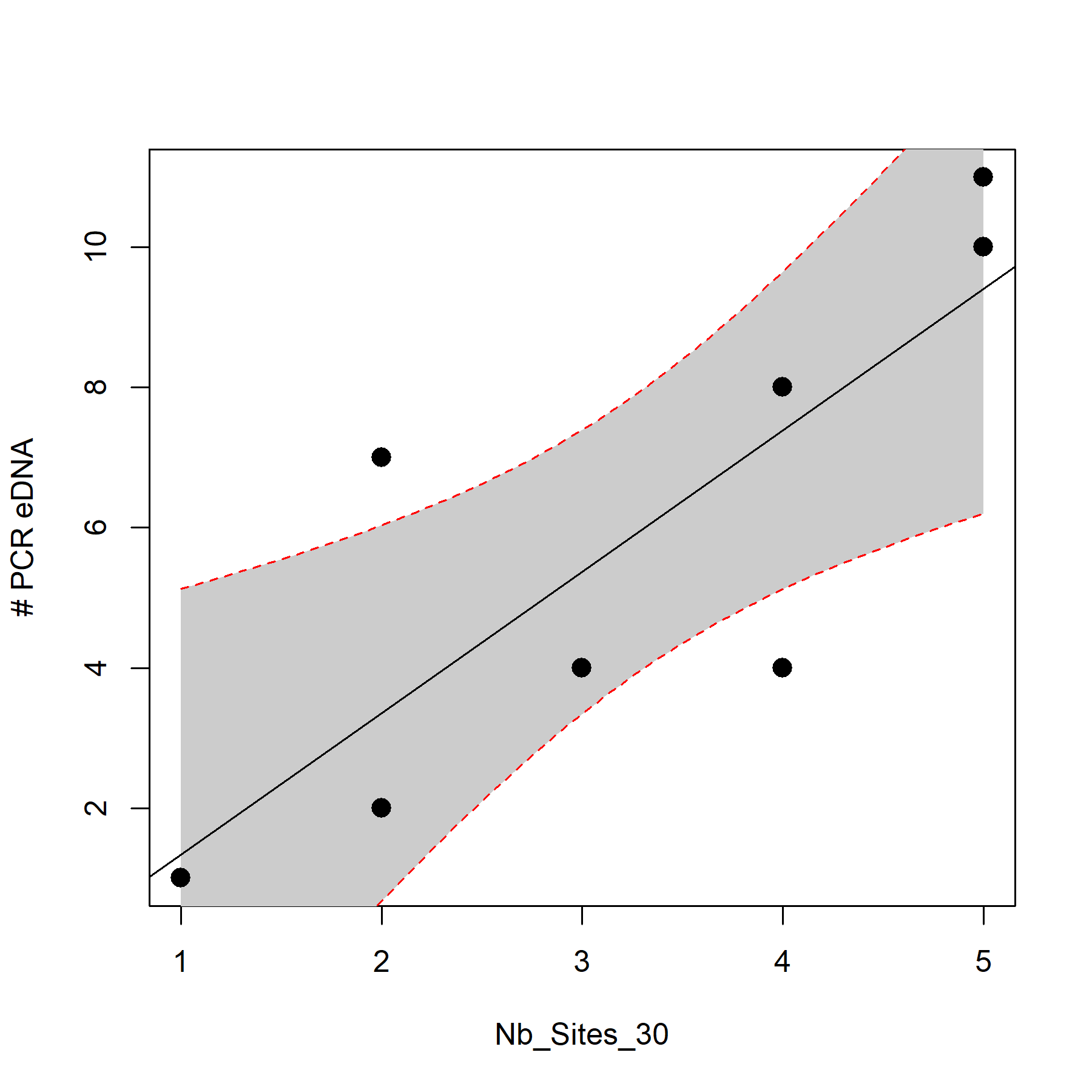

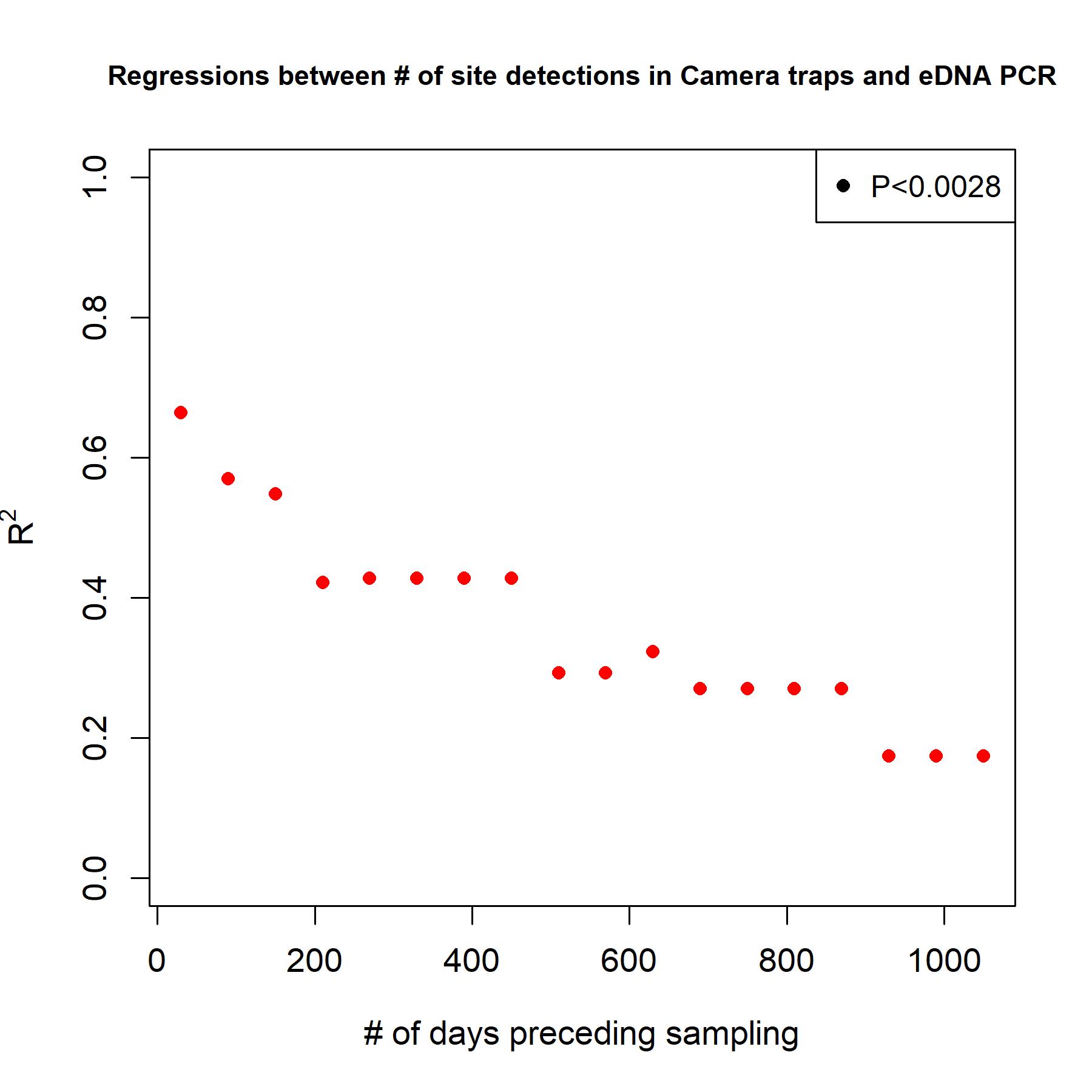

Figure 4 (Top) Quantitative relationship between the number of sites at which species (n=9) are recorded by camera traps and detected in eDNA PCR products (M12s), using a month of camera trapping records preceding sampling. (Bottom) Change in the regression coefficient for the same relationship with increasing camera trap records. Camera trap records from 1 month to 3 years by steps of 60 days. Significant regressions (P<0.05 with Bonferroni correction) are marked with black points.

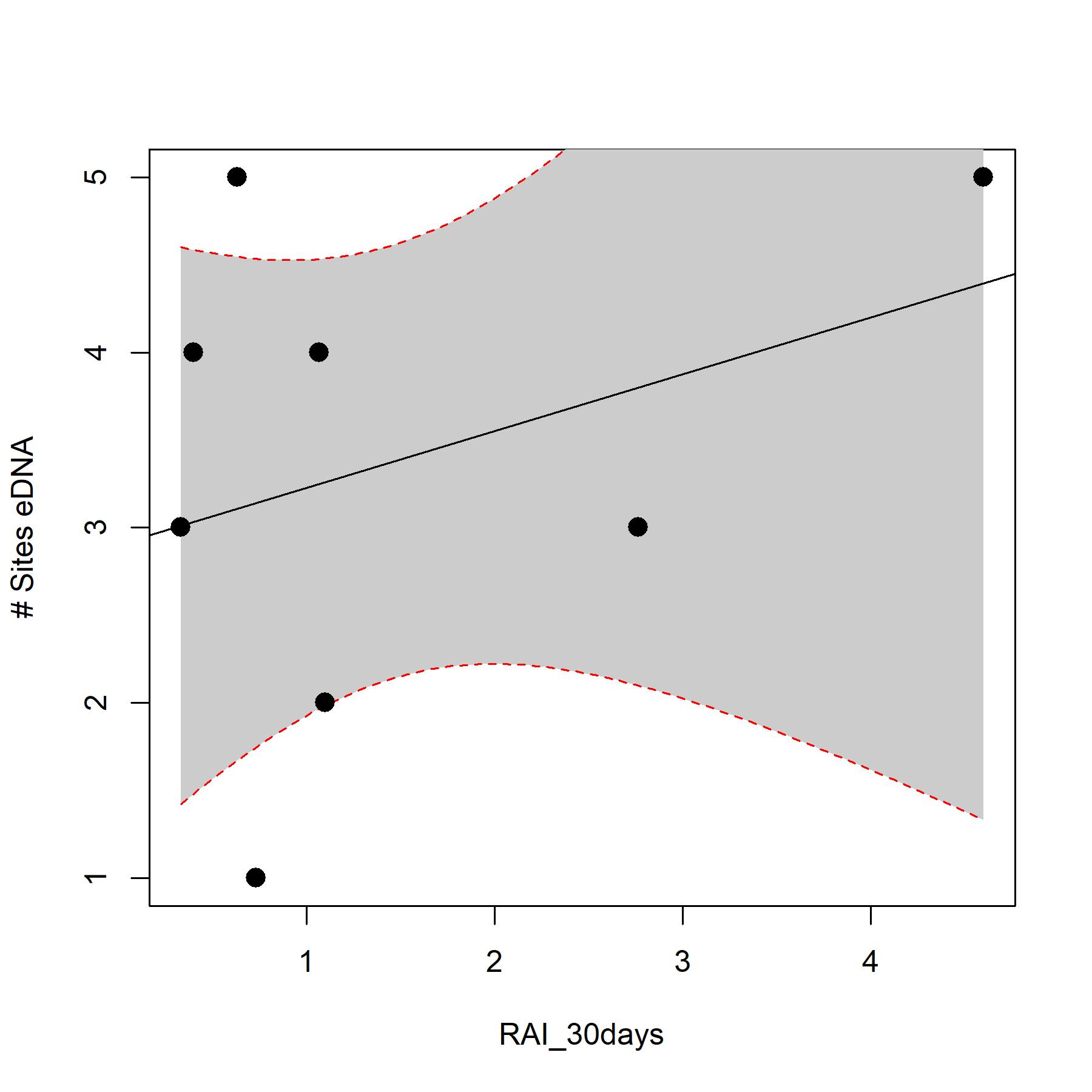

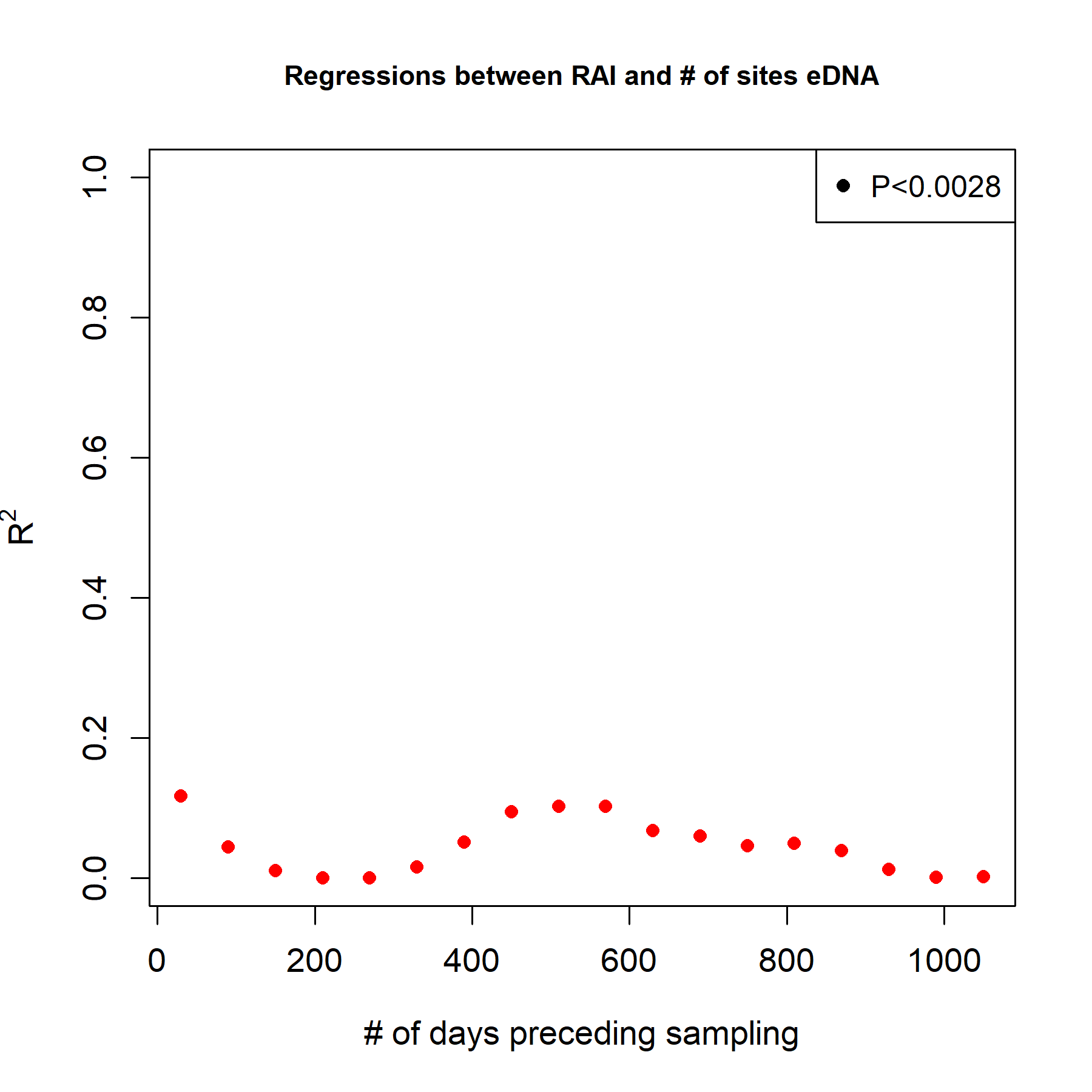

Figure S5 (Top) Quantitative relationship between camera traps RAI (n=9) and the number of sites in eDNA (M12s), using a month of camera trapping records preceding sampling. (Bottom) Change in the regression coefficient for the same relationship with increasing camera trap records. Camera trap records from 1 month to 3 years by steps of 60 days. Significant regressions (P<0.05 with Bonferroni correction) are marked with black points.

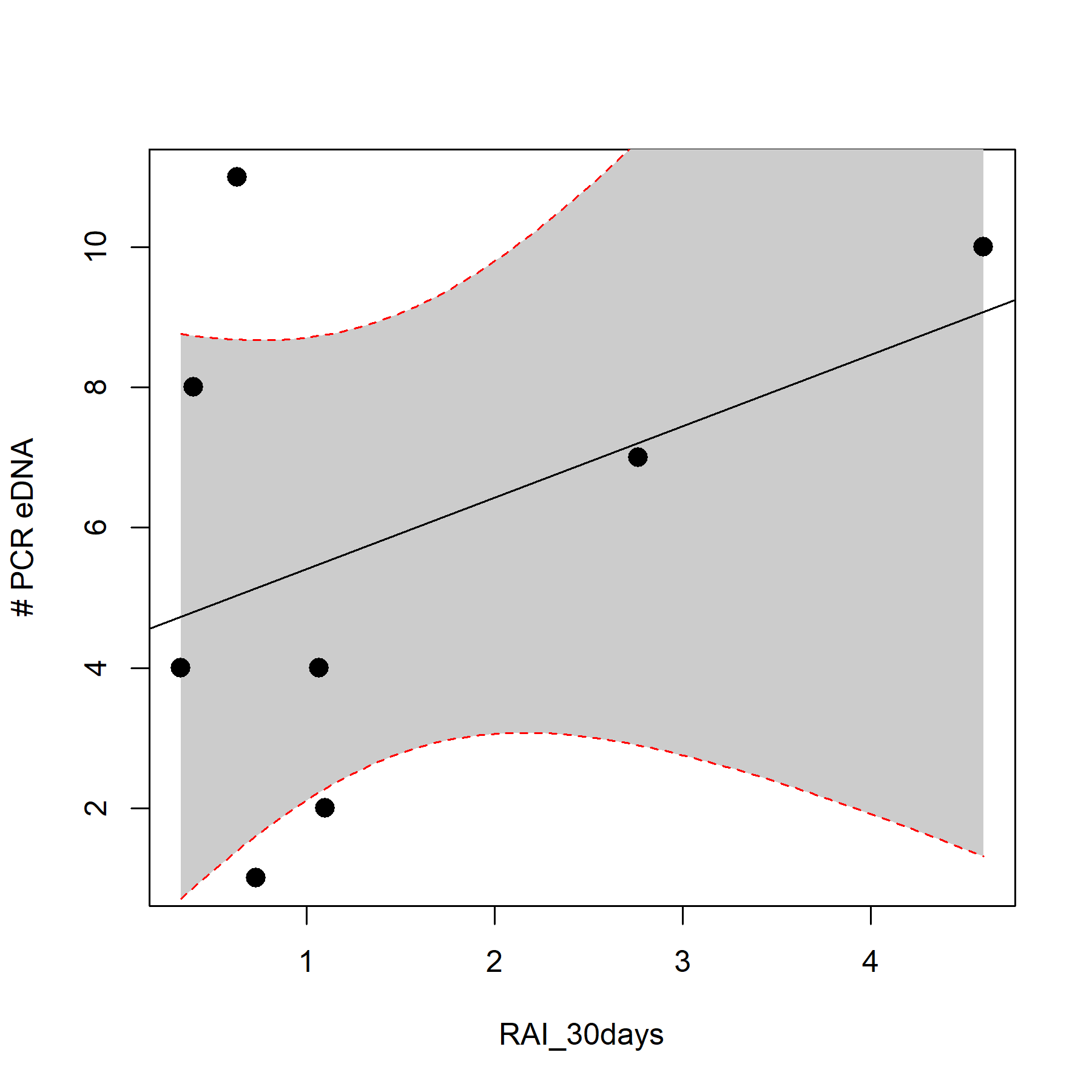

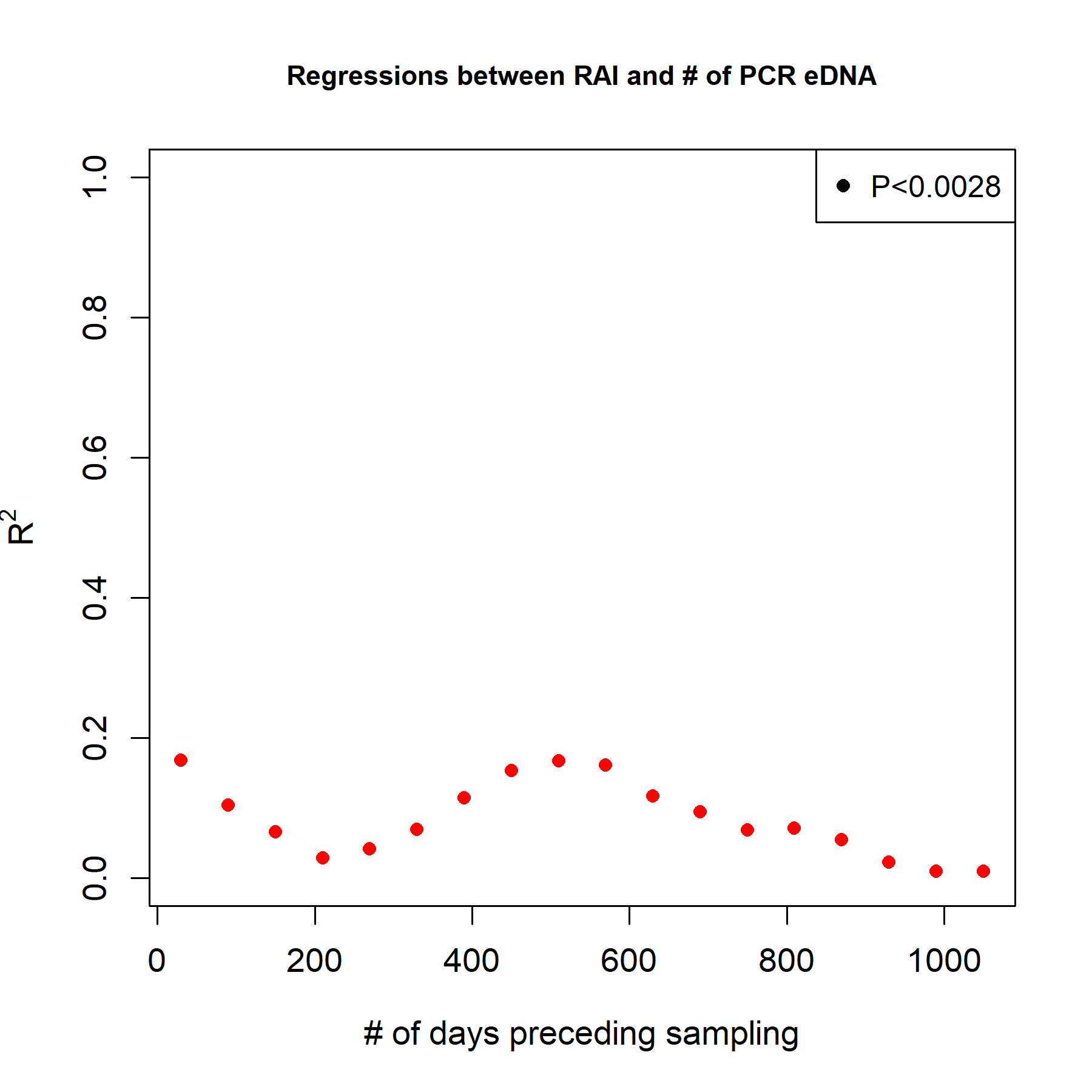

Figure S6 (Top) Quantitative relationship between camera traps RAI (n=9) and the number of detections in eDNA PCR products (M12s), using a month of camera trapping records preceding sampling. (Bottom) Change in the regression coefficient for the same relationship with increasing camera trap records. Camera trap records from 1 month to 3 years by steps of 60 days. Significant regressions (P<0.05 with Bonferroni correction) are marked with black points.

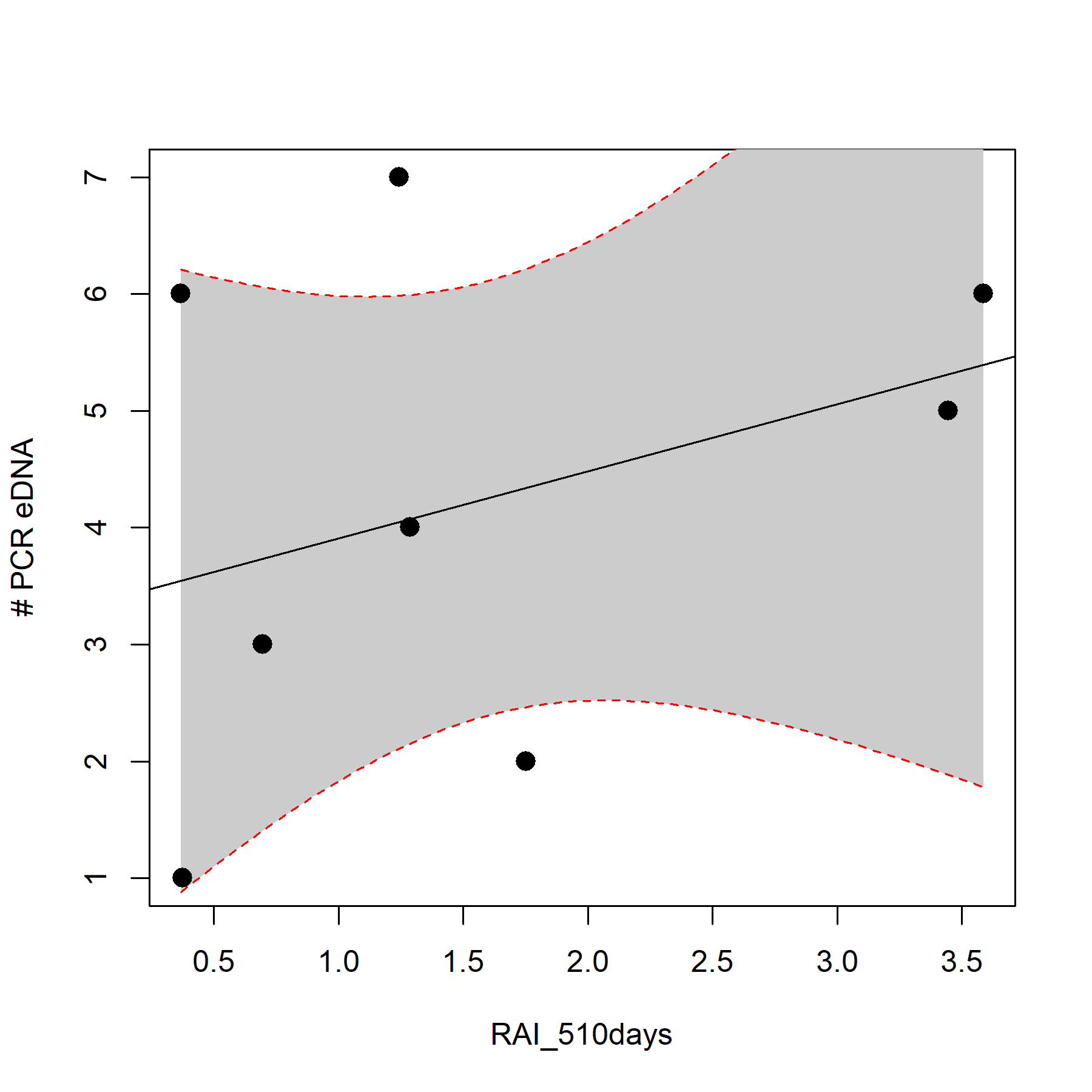

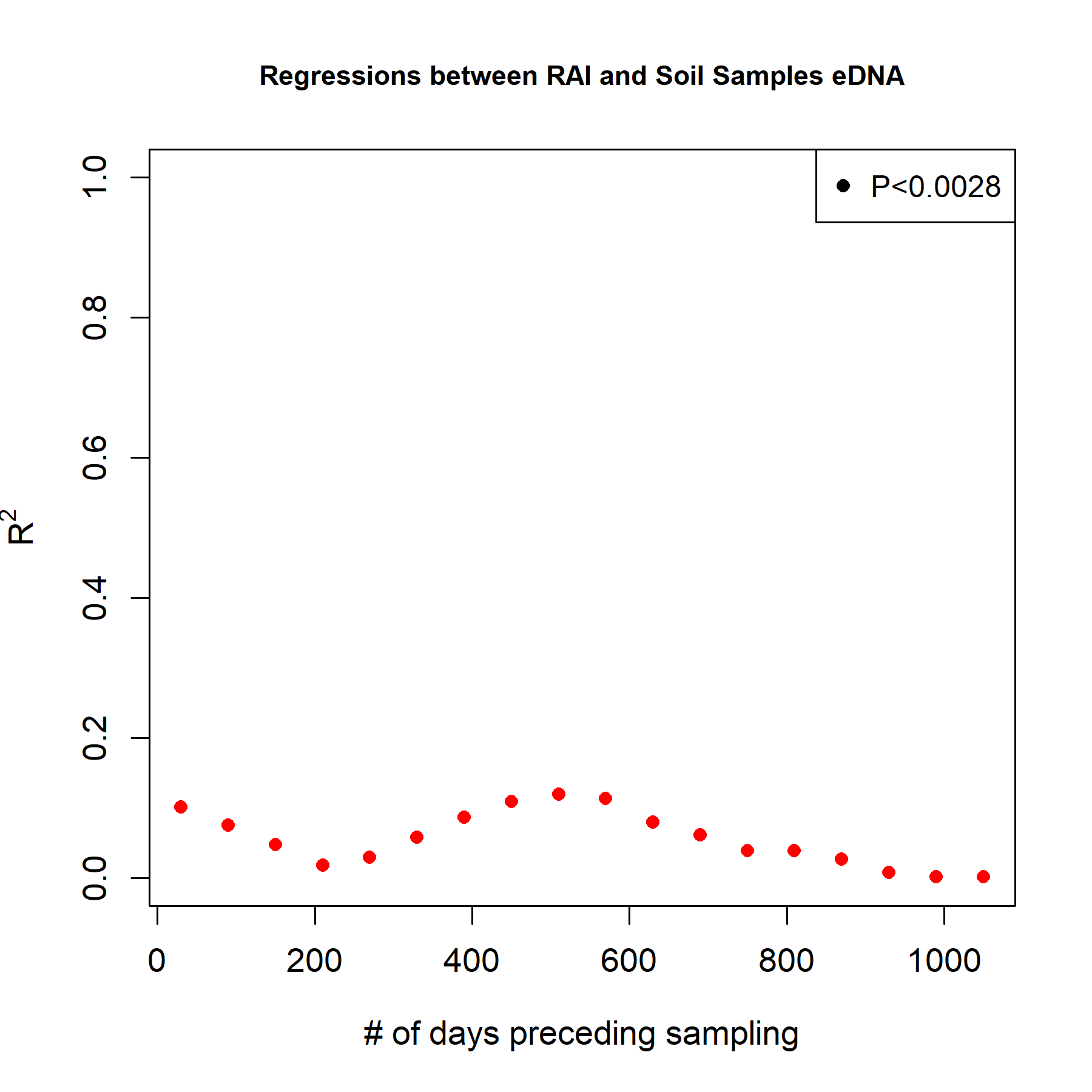

Figure S7 (Top) Quantitative relationship between camera traps RAI (n=9) and the number of detections in eDNA soil samples (M12s), using a 510 days of camera trapping records preceding sampling. (Bottom) Change in the regression coefficient for the same relationship with increasing camera trap records. Camera trap records from 1 month to 3 years by steps of 60 days. Significant regressions (P<0.05 with Bonferroni correction) are marked with black points.

Table S1 Query of the M12s database for all mammals. Shows the number of species queried by order, the number of these found or not found in the M12s database, the percentage of them found in the database, the percentage of those not found in the database but for which sister species at genus or family rank could be found.

| Order name | N | Found | Not found | Genus found | Family found | % found | % found at genus | % found at family |
| --- | --- | --- | --- | --- | --- | --- | --- | --- |
| RODENTIA | 1752 | 432 | 1320 | 679 | 637 | 25% | 51% | 48% |
| CHIROPTERA | 930 | 391 | 539 | 477 | 56 | 42% | 88% | 10% |
| PRIMATES | 374 | 209 | 165 | 136 | 21 | 56% | 82% | 13% |
| EULIPOTYPHLA | 340 | 96 | 244 | 211 | 33 | 28% | 86% | 14% |
| CETARTIODACTYLA | 314 | 231 | 83 | 73 | 8 | 74% | 88% | 10% |
| CARNIVORA | 263 | 136 | 127 | 48 | 79 | 52% | 38% | 62% |
| DIPROTODONTIA | 115 | 70 | 45 | 40 | 5 | 61% | 89% | 11% |
| LAGOMORPHA | 85 | 46 | 39 | 36 | 2 | 54% | 92% | 5% |
| DIDELPHIMORPHIA | 84 | 25 | 59 | 50 | 9 | 30% | 85% | 15% |
| DASYUROMORPHIA | 69 | 64 | 5 | 5 | 0 | 93% | 100% | 0% |
| AFROSORICIDA | 52 | 26 | 26 | 14 | 12 | 50% | 54% | 46% |
| CINGULATA | 20 | 20 | 0 | 0 | 0 | 100% | 0% | 0% |
| SCANDENTIA | 19 | 17 | 2 | 2 | 0 | 89% | 100% | 0% |
| PERAMELEMORPHIA | 18 | 17 | 1 | 1 | 0 | 94% | 100% | 0% |
| MACROSCELIDEA | 17 | 14 | 3 | 2 | 1 | 82% | 67% | 33% |
| PERISSODACTYLA | 16 | 14 | 2 | 1 | 0 | 88% | 50% | 0% |
| PILOSA | 10 | 10 | 0 | 0 | 0 | 100% | 0% | 0% |
| PAUCITUBERCULATA | 6 | 2 | 4 | 3 | 1 | 33% | 75% | 25% |
| PHOLIDOTA | 6 | 4 | 2 | 2 | 0 | 67% | 100% | 0% |
| SIRENIA | 5 | 2 | 3 | 2 | 1 | 40% | 67% | 33% |
| MONOTREMATA | 5 | 3 | 2 | 2 | 0 | 60% | 100% | 0% |
| HYRACOIDEA | 4 | 2 | 2 | 1 | 1 | 50% | 50% | 50% |
| PROBOSCIDEA | 2 | 2 | 0 | 0 | 0 | 100% | 0% | 0% |
| DERMOPTERA | 2 | 1 | 1 | 0 | 1 | 50% | 0% | 100% |
| NOTORYCTEMORPHIA | 2 | 2 | 0 | 0 | 0 | 100% | 0% | 0% |
| TUBULIDENTATA | 1 | 1 | 0 | 0 | 0 | 100% | 0% | 0% |
| MICROBIOTHERIA | 1 | 1 | 0 | 0 | 0 | 100% | 0% | 0% |
| **Total** | **4512** | **1838** | **2674** | **1785** | **867** | **41%** | **67%** | **32%** |
